# Acquired resistance to immune checkpoint inhibitors is associated with hypoxia and ECM remodeling in colorectal cancer

**DOI:** 10.64898/2026.04.27.720591

**Authors:** Astrid Zedlitz Johansen, Hannes Linder, Cecilie Ølvang Madsen, Kevin James Baker, Klaire Yixin Fjæstad, Shawez Khan, Nadia Kolvig Czajkowski, Christian Thygesen, Majken Siersbæk, Frederik Otzen Bagger, Marco Donia, Lars Henning Engelholm, Lars Grøntved, Daniel Hargbøl Madsen

**Author notes:** Correspondence: Daniel Hargbøl Madsen, National Center for Cancer Immune Therapy (CCIT-DK), Department of Oncology, Copenhagen University Hospital - Herlev and Gentofte, DK-2730 Herlev, Denmark.

## Abstract

Acquired resistance to immune checkpoint inhibitors (ICIs) limits the durability of therapeutic responses across multiple cancer types, yet the underlying mechanisms remain poorly defined. Using the murine MC38 colorectal cancer model, we established an *in vivo* model recapitulating clinical response patterns, including complete regression, primary resistance, and acquired resistance. Tumors with acquired resistance progressed after initial benefit from combined anti-PD-1 and anti-CTLA-4 therapy and maintained resistance upon retransplantation into naïve hosts, indicating a cancer cell-intrinsic driver of resistance. Whole-genome sequencing showed no mutations in antigen presentation or IFN-γ signaling pathways previously described in single clinical cases of acquired resistance, and functional assays confirmed preserved antigenicity and IFN-γ responsiveness. Transcriptomic and metabolic profiling of resistant cancer cells instead revealed metabolic reprogramming characterized by enhanced mitochondrial respiration and enrichment of hypoxia-related gene signatures, suggesting cancer cell–intrinsic adaptations that reshape the tumor microenvironment. Tumors with acquired resistance exhibited an increase in tumor-associated macrophages, and these macrophages displayed enriched transcriptional signatures of hypoxia, angiogenesis, and extracellular matrix (ECM) remodeling. Proteomic analysis of ECM-enriched tumor fragments showed accumulation of proteoglycans and enzymes such as lysyl oxidase, consistent with active matrix-remodeling. These changes coincided with altered T cell activation characterized by reduced cytotoxic gene expression and decreased CD44 expression on tumor-infiltrating T cells, suggesting impaired effector functions within the resistant microenvironment. This study identifies non-genetic mechanisms of acquired resistance to ICIs and highlights metabolic and ECM remodeling programs as promising therapeutic targets to prevent or reverse acquired resistance to ICIs.

## INTRODUCTION

Immune checkpoint inhibitors (ICIs) have revolutionized cancer therapy, transforming outcomes for patients with many different types of cancer. By targeting immune regulatory pathways such as PD-1/PD-L1 and CTLA-4, ICIs release robust anti-tumor immune responses that lead to unprecedented survival benefits (1,2). Despite their transformative potential, two major obstacles often hinder the clinical success of ICIs: primary resistance, where tumors fail to respond to treatment from the beginning; and acquired resistance (also referred to as secondary resistance), where initially responsive tumors relapse during therapy. Acquired resistance poses a major challenge in solid tumors, affecting up to 65% of patients who initially benefit from ICIs and limiting the durability of therapeutic responses (3–5).

Unlike primary resistance, acquired resistance emerges after an initial therapeutic response, warranting its study as a distinct entity (6). Factors involved in acquired resistance may either be pre-existing at baseline, acquired genetically, or acquired through non-genetic adaptation (7). Early studies of single clinical cases with acquired resistance primarily described genomic alterations, such as inactivating mutations in two pathways: MHC-I antigen presentation and interferon-gamma (IFN-γ) signaling (8–10). These mutations reduce the cancer cells’ ability to present antigens and respond to IFN-γ, thereby enabling them to evade CD8^+^ T cell-mediated immune killing. While these findings provided initial clues about the genetic landscape of acquired resistance, later studies uncovered significant heterogeneity in acquired resistance mechanisms, highlighting that the underlying molecular cause of resistance remains unknown for most patients (11–13).

Recent insights into acquired resistance to ICIs suggest a complex interplay of tumor-intrinsic and extrinsic mechanisms and a potentially prominent role of non-genomic factors (14). Recognizing the challenges of studying acquired resistance in patients due to high interpatient variability and limited sample availability at time of resistance, syngeneic mouse models have demonstrated their utility as controlled systems to provide insight into the biology of acquired resistance (12,15–20). Studies using these models typically involve repeated implantation of cancer cells into immunocompetent hosts under continued ICI treatment, thereby exerting selective pressure that enriches for therapy-resistant cells. However, this methodology may introduce a selection bias toward tumor-intrinsic mechanisms, such as dysfunctional IFN-γ signaling (12). This raises the question of whether current preclinical models recapitulate the complexity of acquired resistance as it occurs in patients.

To address this, we demonstrate that ICI treatment in a mouse model of colorectal cancer can lead to heterogeneous therapeutic responses upon initial treatment, which recapitulate clinical response patterns. Therapeutic resistance is driven by the adaptation of tumor cells to ICI-induced immune pressure, which results in a shift towards a metabolic state characterized by enhanced mitochondrial respiration. This metabolic reprogramming fosters hypoxic conditions within the tumor and activates hypoxia-associated signaling pathways. Furthermore, these changes contribute to tumor progression by remodeling the tumor microenvironment (TME), particularly through alterations in the extracellular matrix (ECM). These changes are accompanied by phenotypic and functional alterations in the tumor-infiltrating T cells, suggesting diminished effector activity within the remodeled TME.

## RESULTS

### Modelling acquired resistance to ICIs in a mouse model of colorectal cancer

The MC38 colon cancer cell line was used to establish a syngeneic mouse model of acquired resistance as it is sensitive to treatment with ICIs (21,22). MC38 cancer cells were subcutaneously inoculated in the flank of immunodeficient *Rag2* KO or wild-type C57BL/6 mice. When the tumor volume reached 50 mm^3^, wild-type mice were randomized into a control or ICI treatment group (Figure 1A). Tumors in *Rag2* KO mice progressed faster than tumors in wild-type mice, leading to a shorter survival of these mice (Figures 1B and 1C), indicating that the tumors were susceptible to immune-mediated control. Treatment with ICIs delayed tumor growth, and in 3 out of 15 mice, the tumors were cleared (Figure 1B). The experiment was ended when tumors reached a pre-defined size (Supplementary Figure 1A), which led to a prolonged survival of ICI-treated mice compared to control-treated mice (Figure 1C).

**Figure 1.**
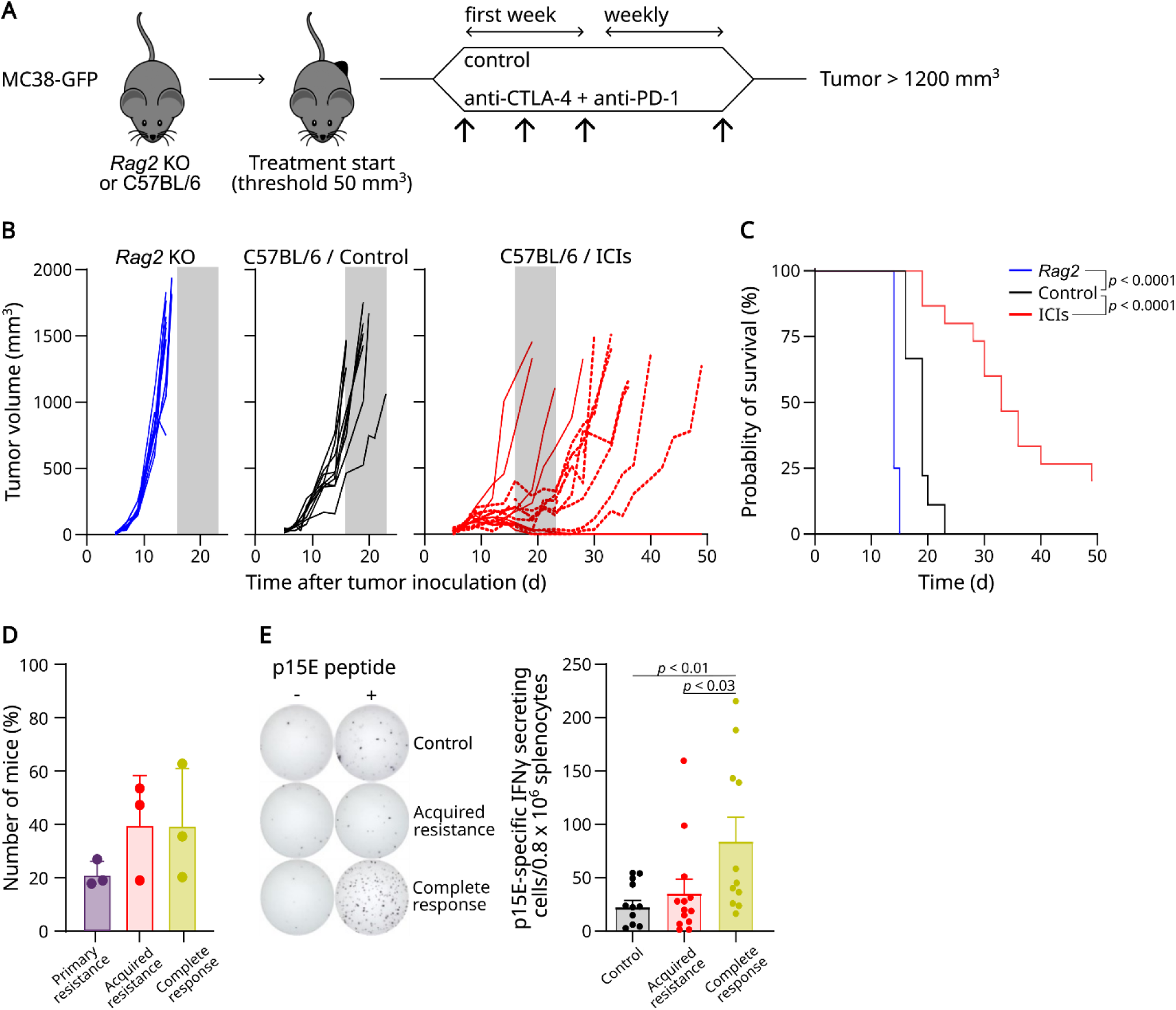
Heterogenous responses to immune checkpoint inhibitors in a mouse model of colorectal cancer. (A) MC38 cells were injected into the flank of *Rag2* KO or wild-type C57BL/6 mice subjected to control or ICI treatment initiated at tumor volumes > 50 mm^3^. (B) Individual tumor growth curves. Grey shaded area indicates the time window at which control tumors reached endpoint. Dotted lines highlight tumors with acquired resistance. (C) Survival curves for *Rag2* KO mice (n = 8), control-treated (n = 9), and ICI-treated wildtype C57BL/6 mice (n = 15). (D) Response patterns in the ICI-treated group across three independent experiments. (E) IFN-γ ELISpot responses. Statistical significance was assessed using the log-rank test (C) and the Mann-Whitney test (E). Bars represent means and error bars represent standard deviation (SD).

Notably, heterogeneous responses to ICIs were observed in the ICI-treated group. Acquired resistance was defined as an initial delay in tumor growth followed by tumor progression, leading to a survival > 50% longer than the average survival time for control-treated mice. Applying this definition to the ICI-treated group, 4 (27%) mice displayed primary resistance, 8 (53%) mice displayed acquired resistance, and 3 (20%) mice had complete regression of their tumors (Figure 1B). These rates were comparable across three independent experiments (Figure 1D and Supplementary Figure 1).

To investigate whether acquired resistance was associated with decreased anti-tumor T cell reactivity, splenocytes from control- and ICI-treated mice were collected and tested for their reactivity against the p15E peptide, an MHC class I-restricted cancer T cell epitope present in the MC38 model (23). While the number of splenocytes secreting IFN-γ was comparable between control- and ICI-treated mice with acquired resistance, mice with complete tumor regression demonstrated a higher frequency of tumor-specific T cells (Figure 1E). These data suggest that the risk of acquired resistance is associated with reduced expansion or maintenance of tumor-specific T cells, while stronger T cell responses are linked to more durable tumor control.

### Cancer-intrinsic changes drive acquired resistance

To determine whether the acquired resistance response pattern reflected a stable cancer cell-intrinsic phenotype or required further *in vivo* selection, MC38 tumors were harvested at terminal volume and transplanted into recipient mice either as tumor fragments or as sorted cancer cells, followed by ICI treatment (Figure 2A). Tumors derived from cancer cells of ICI-naïve control tumors remained sensitive to ICI (Figure 2B), whereas those derived from cancer cells of tumors with acquired resistance were refractory to the treatment (Figure 2C). This demonstrates that acquired resistance is driven by a cancer cell-intrinsic phenotype that does not require continued *in vivo* selection. Unexpectedly, transplantation of tumor fragments yielded a different outcome. Tumors from control and acquired resistance fragments both displayed primary resistance to ICI treatment (Figure 2D-E). This suggests that factors within the TME can contribute to resistance independently of prior ICI treatment. Interestingly, similar primary resistance was not observed when the fragments originated from tumors formed in immunodeficient *Rag2* KO mice (Figure 2F). Together, these data indicate that ICI resistance exists at two levels: a cancer cell-intrinsic change and an immune-mediated conditioning of the TME. Even when cancer cells themselves remain sensitive to ICI treatment as observed with cancer cells from control tumors, the immune system exerts a selective pressure that conditions tumors toward primary resistance when the intact TME is transferred (Figure 2G).

**Figure 2.**
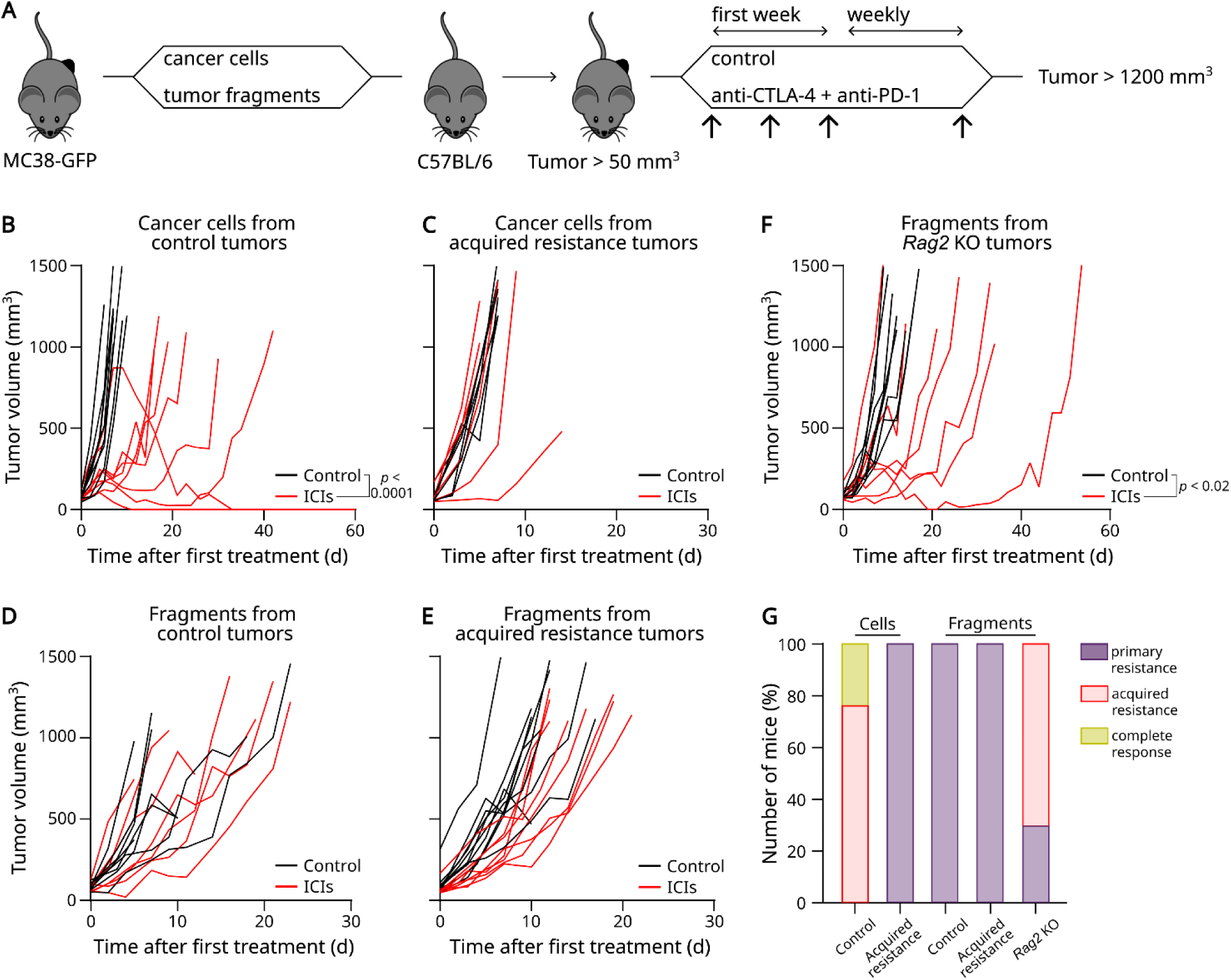
Transplantation of acquired resistance tumors leads to primary resistance. (A) Cancer cells and tumor fragments from control-treated, ICI-treated with acquired resistance, and *Rag2* KO mice were grafted into recipient, tumor-naïve mice subjected to control or ICI treatment initiated at tumor volumes > 50 mm^3^. (B-C) Individual tumor growth curves for tumors derived from cancer cells from control (B) and acquired resistance tumors (C). (D-E) Individual tumor growth curves for tumors derived from implanted fragments from control (D) and acquired resistance tumors (E). (F) Individual tumor growth curves for tumors derived from implanted fragments from *Rag2* KO tumors. (G) Distribution of response patterns in recipient mice. Tumor growth differences were analyzed using a linear mixed-effects model (B-F).

### Acquired resistance occurs without genetic loss of antigenicity or IFN-γ responsiveness

To investigate if genomic alterations were associated with acquired resistance, whole-genome sequencing was performed on DNA isolated from the parental MC38 cell line and cancer cells sorted from control and acquired resistance tumors. The MC38 cell line is characterized by a high mutational burden and sensitivity to ICIs, including endogenous CD8⁺ T cell responses against neoantigens. While lacking common colorectal cancer mutations such as *KRAS* and *APC*, MC38 harbors heterogeneous *Trp53* mutations and previously described missense mutations and neoantigens (Supplementary Table 1) (21,24,25). Despite the absence of canonical mismatch repair or polymerase mutations, MC38 exhibits a high mutational load (∼87 mutations/Mb), supporting its relevance as a preclinical model of hypermutated, ICI-responsive tumors. The overall somatic mutational burden remained unchanged across all samples, with a median of 2,587 mutations (Figure 3A). The majority of single-nucleotide variants were transversions, particularly C>A/G>T substitutions, but there were no differences between the samples (Figure 3B). To explore whether acquired resistance involved reduced tumor antigenicity, the number of HLA class I-restricted neoantigens was predicted *in silico*. No decrease in the number of neoantigens was observed (Figure 3C).

**Figure 3.**
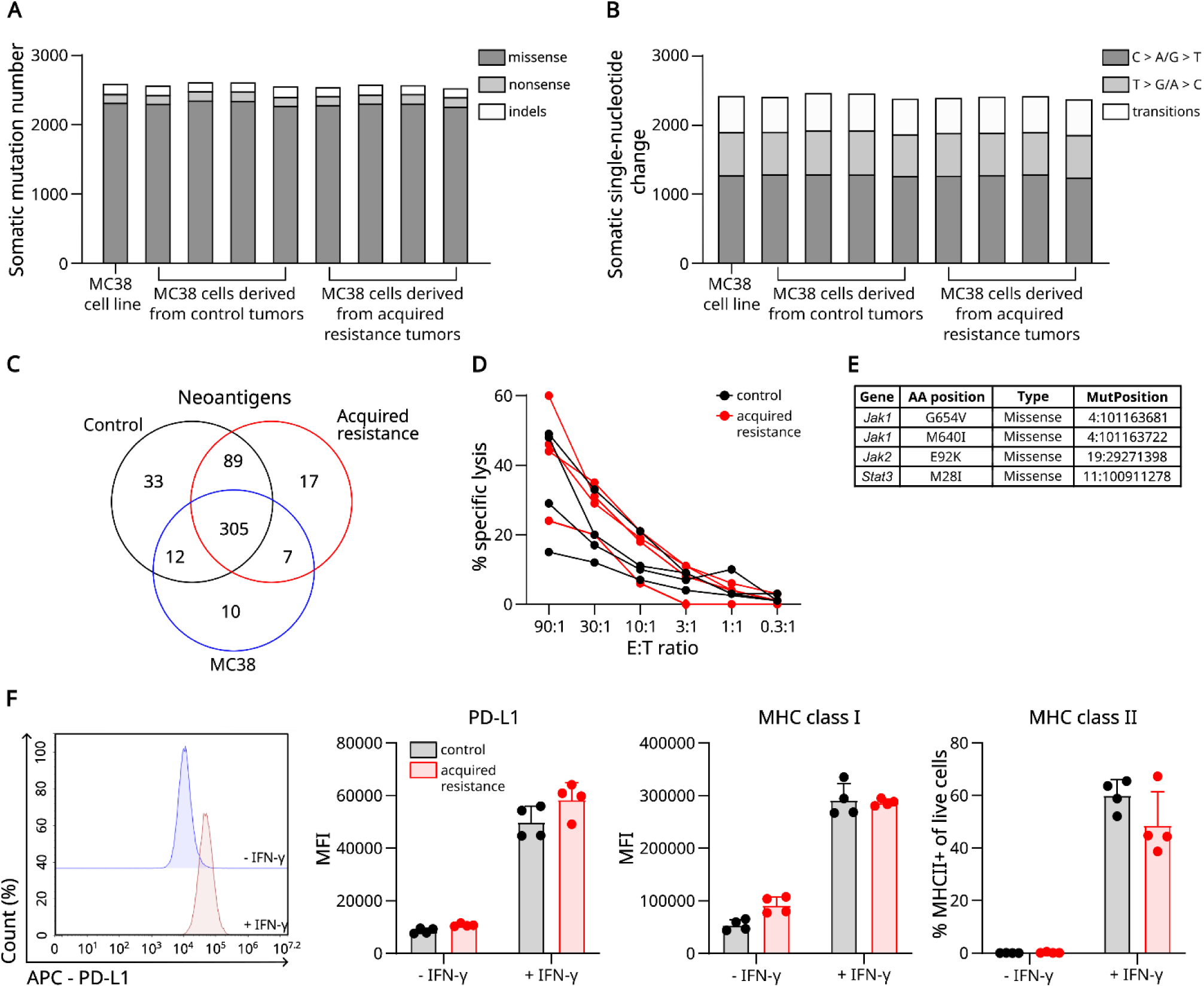
Genomic landscape of acquired resistance in the MC38 model. (A) Somatic mutation counts by type. (B) Distribution of somatic single-nucleotide variants. (C) *In silico* predicted HLA class-I restricted neoantigen counts. (D) Specific lysis percentages of cancer cell lines derived from control and acquired resistance tumors (E:T; effector-to-target ratio). (E) Overview of somatic mutations in the IFN-γ signaling pathway. (F) Representative histograms showing PD-L1 expression in cultured cancer cells ± IFN-γ stimulation (left) and bar graphs quantifying surface expression of PD-L1, MHC class I, and MHC class II (right). Bars represent means and error bars represent SD.

Given that mutations affecting antigenicity and IFN-γ responsiveness have been reported in patients with melanoma and lung cancer developing acquired resistance to ICIs, similar mechanisms were investigated. No mutations were detected in genes associated with antigen processing and presentation including *B2m*, *H2-D1*, *H2-K1*, *Tap1*, and *Tap2* (Supplementary Table 2) . Antigen presentation was further assessed functionally by co-culturing OT-1 T cells with cancer cells pulsed with the SIINFEKL peptide, which is recognized by OT-1 T cells. T cell-mediated killing was comparable between control and ICI-resistant cell lines (Figure 3D), and no killing occurred in the absence of peptide loading (Supplementary Figure 2B), demonstrating intact antigen presentation machinery and similar susceptibility to T cell-mediated killing. Mutations in *Jak1*, *Jak2*, and *Stat3* were identified in all samples, including the parental cell line, indicating that these mutations were present prior to ICI treatment and were not specifically acquired during the development of resistance (Figure 3E). Nevertheless, to assess whether the mutations affected IFN-γ signaling at a functional level, cancer cell lines derived from control and acquired resistance tumors were exposed to IFN-γ. IFN-γ stimulation increased PD-L1, MHC class I, and MHC class II expression to a similar extent in both groups (Figure 3F), indicating preserved IFN-γ responsiveness. Together, these results suggest that genetic resistance mechanisms, including impaired antigen presentation and defective IFN-γ signaling, do not drive acquired resistance in this model.

### Hypoxia signaling and metabolic cancer cell rewiring characterize acquired resistance

To identify alternative biological pathways related to acquired resistance, cancer cells were sorted from control and acquired resistance tumors. Following RNA isolation and sequencing, differential gene expression analysis was performed. The transcriptome distinctly differed between cancer cells from control and acquired resistance tumors (Supplementary Figure 2C). A total of 867 genes were upregulated in acquired resistance cancer cells, while 437 were downregulated (Figure 4A). Consistent with the absence of genomic alterations in genes related to antigen processing, presentation, or IFN-γ responsiveness, transcript levels of these pathways remained unaltered between control and acquired resistance cancer cells (Supplementary Figure 2D). Enrichment analysis of gene ontology terms highlighted that terms related to leukocyte-mediated immunity and ECM were upregulated in acquired resistance (Figure 4B). Interestingly, gene set enrichment analysis (GSEA) revealed that hypoxia was associated with acquired resistance (Figure 4C and Supplementary Figure 2E). TNF-α signaling via NF-κB and epithelial-mesenchymal transition were also shown to be associated with acquired resistance (Figure 4C and Supplementary Figure 2E).

**Figure 4.**
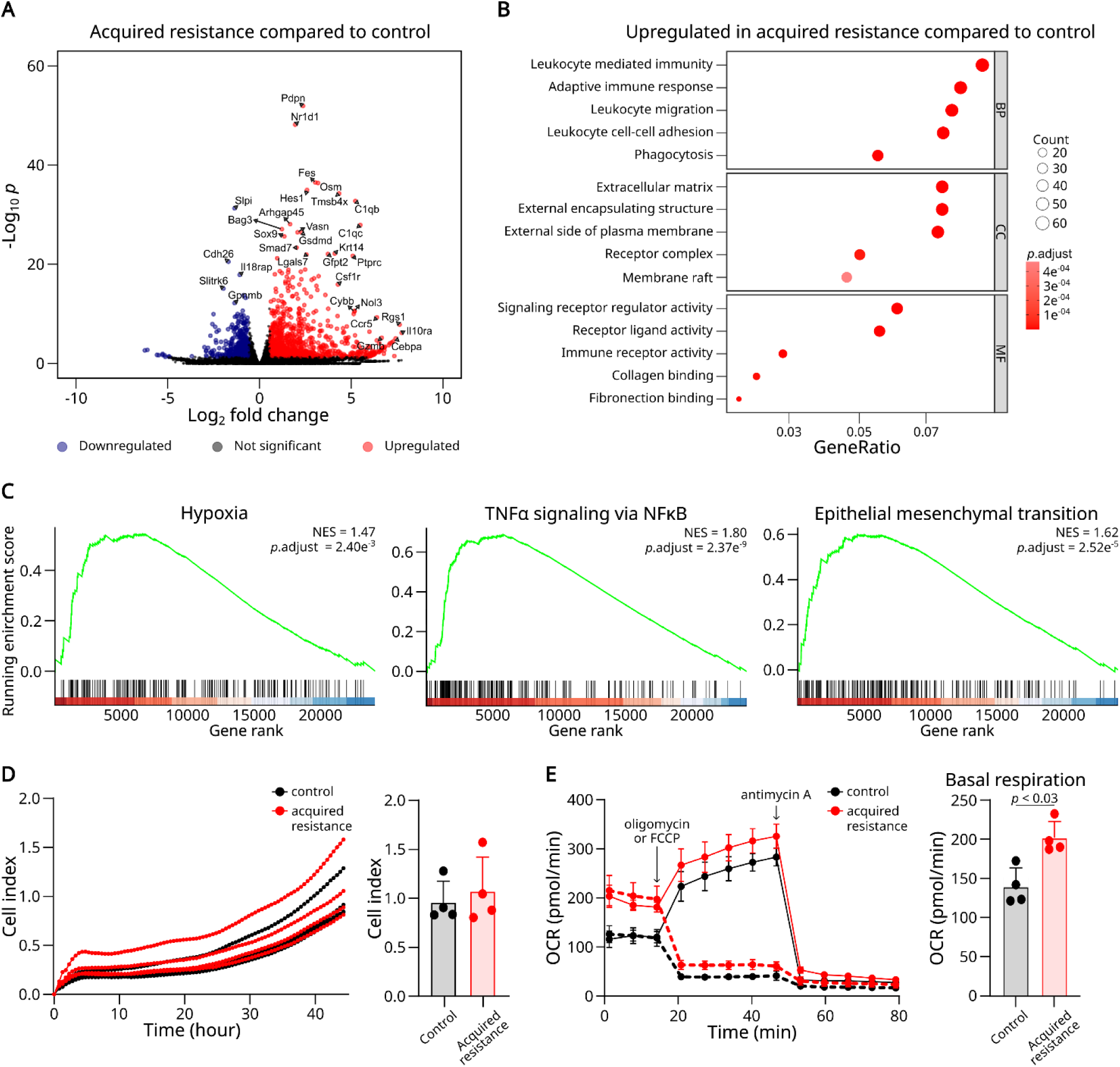
Transcriptomic and metabolic adaptations in resistant cancer cells. (A) Differentially expressed genes in cancer cells sorted from acquired resistance tumors (n = 2) versus control tumors (n = 4). (B) Upregulated gene ontology terms for biological processes (BP), cellular components (CC), and molecular functions (MF) in acquired resistance. (C) Enrichment plots of top-scoring gene sets. (D-E) To further characterize resistant cancer cells, cancer cell lines were established from control and acquired resistance tumors (n = 4 per group). (D) Proliferative capacities over time and cell index at 44 hours. (E) Oxygen consumption rates (OCR) of representative cell lines and a bar graph of basal OCR. Statistical comparisons were performed using a Wald test (A), hypergeometric test (B) or permutation-based multiple testing procedure (C) with Benjamini-Hochberg correction (A-C) and the Mann-Whitney test (E). Bars represent means and error bars represent SD.

Tumor hypoxia can develop due to altered metabolism or increased cell proliferation. Cancer cell proliferation *ex vivo* was similar between the two groups (Figure 4D), but assessment of mitochondrial respiration revealed higher basal oxygen consumption rates in cancer cells with acquired resistance (Figure 4E). This result showed that acquired resistance is associated with a metabolic shift of the cancer cells characterized by enhanced mitochondrial respiration. Further analysis showed that ATP turnover was similar and that control cancer cell lines exhibited a higher spare respiratory capacity (Supplementary Figure 2F), suggesting that the elevated baseline oxygen consumption rate was not attributable to differences in mitochondrial mass.

These findings suggest that acquired resistance to ICI treatment is driven by significant transcriptomic changes, including upregulation of immune-related pathways, ECM remodeling, and metabolic reprogramming. The enrichment of a hypoxia signature and increased basal oxygen consumption in resistant cancer cells point to enhanced metabolic activity, which may promote immune evasion.

### Hypoxia-associated TAM reprogramming and altered T cell activation in acquired resistance

To investigate the tumor immune microenvironment in the context of acquired resistance, tumors were analyzed using flow cytometry. The overall composition of major cell classes differed between control and acquired resistance tumors. The percentage of cancer cells was lower (*p* < 0.05) and leukocytes showed a trend towards an increase (*p* = 0.1) in resistant tumors (Figure 5A).

**Figure 5.**
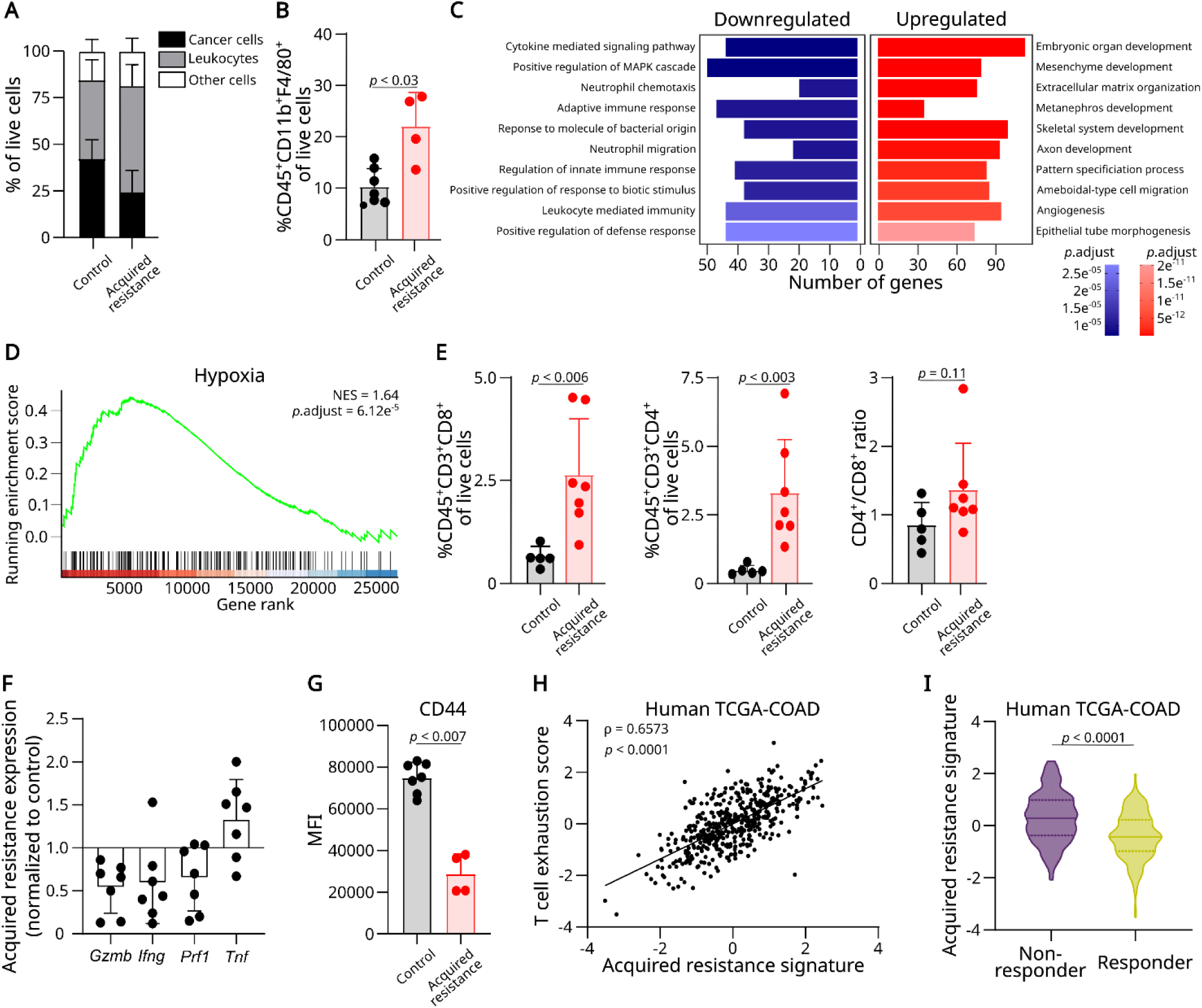
Tumor immune microenvironment in acquired resistance. (A) Tumor composition analyzed at the time of terminal tumor volume by flow cytometry (n = 7 control; n = 4 acquired resistance). (B) TAM abundance. (C-D) TAMs were sorted from control (n = 4) and acquired resistance tumors (n = 2). (C) Gene ontology analysis of differentially expressed genes in TAMs from acquired resistance tumors versus control. (D) Enrichment plot of the hypoxia gene set. (E) T cell abundance. (F) Expression of genes encoding cytotoxic molecules from two independent *in vivo* experiments (n = 3-4 per group per experiment), shown as RNA-seq counts normalized to average control expression. (G) CD44 expression in CD8^+^ T cells. (H) Correlation between the acquired resistance signature score and CD8⁺ T cell exhaustion score. Each point represents an individual sample in the TCGA-COAD cohort. (I) Association between predicted ICI responder status and the acquired resistance signature score in the TCGA-COAD cohort (distribution with first quartile, median, and third quartile). Statistical comparisons were performed using the Mann-Whitney test (A-B, E, G, I), a hypergeometric test (C) or permutation-based multiple testing procedure (D) with Benjamini-Hochberg correction (C-D), and Spearman correlation (H). Bars represent means and error bars represent SD.

By comparison of specific immune cell subsets, the trend toward more leukocytes in tumors with acquired resistance was primarily due to an increase in tumor-associated macrophages (TAMs) (Figure 5B). Cancer cells from acquired resistance tumors expressed higher levels of several chemokines implicated in monocyte and macrophage recruitment, including *Ccl2*, *Ccl3*, *Ccl7*, and *Cxcl14* (Supplementary Figure 3A). Other myeloid cell populations, including monocytic and polymorphonuclear myeloid-derived suppressor cells (MDSCs), did not change in number (Supplementary Figure 3B). TAMs are a diverse cell population, which can exert both pro- and anti-tumorigenic effects. To further assess their phenotype, TAMs were sorted from control and acquired resistance tumors (Supplementary Figure 3C), followed by RNA isolation and sequencing. Differential gene analysis showed that the transcriptome clearly distinguished between TAMs from ICI-naïve control tumors and ICI-treated tumors with acquired resistance (Supplementary Figure 3D). A total of 1429 genes were upregulated in TAMs from acquired resistance tumors, while 1302 were downregulated. Enrichment analysis of gene ontology terms highlighted terms related to ECM regulation and angiogenesis as upregulated and immune functions as downregulated in TAMs from acquired resistance tumors (Figure 5C). GSEA revealed that hypoxia was again associated with acquired resistance (Figure 5D and Supplementary Figure 3E).

Additionally, there was an increased number of CD8⁺ and CD4⁺ T cells in acquired resistance tumors (Figure 5E). Despite increased T cell infiltration, the relative expression of cytotoxic effector genes (*Gzmb*, *Ifng*, and *Prf1*) tended to be lower in acquired resistance tumors, whereas *Tnf* expression showed an opposite trend (Figure 5F). CD8⁺ T cells exhibited decreased CD44 expression in acquired resistance tumors (Figure 5G), indicating an altered activation status. Taken together, these findings suggest that although T cells accumulate in acquired resistance tumors, their cytotoxic function was impaired. To examine whether the observed link between reduced T cell cytotoxicity and changes in cancer cells and TAMs from resistant tumors were also observed in patients, an acquired resistance signature was generated by intersecting shared leading-edge genes from hypoxia and EMT pathways (Supplementary Table 3). The signature was compared with the T cell exhaustion score and ICI response status defined by the TIDE framework (26). In the TCGA-COAD cohort, this signature positively correlated with CD8⁺ T cell exhaustion (Figure 5H). Moreover, higher signature scores were observed in patients with cancer classified as non-responders (Figure 5I), linking hypoxia and EMT-driven remodeling in acquired resistance to T cell dysfunction and therapeutic resistance.

### ECM remodeling in resistant tumors

Signatures indicative of hypoxia and ECM remodeling were enriched in cancer cells and TAMs isolated from acquired resistance tumors. As hypoxia can promote abnormal angiogenesis (27), RNA was isolated from whole tumors and sequenced. Importantly, the master hypoxia regulator *HIF-1α* was upregulated in acquired resistance tumors (*p* < 0.0001) (Figure 6A). In addition, several hypoxia-associated pro-angiogenic factors were upregulated in acquired resistance tumors (Figure 6A). Paraffin-embedded tissue sections from control and acquired resistance tumors were also immuno-stained for the endothelial marker CD34. A tendency toward an increased CD34^+^ area was observed in acquired resistance tumors (Figure 6B), consistent with enhanced vascular remodeling.

**Figure 6.**
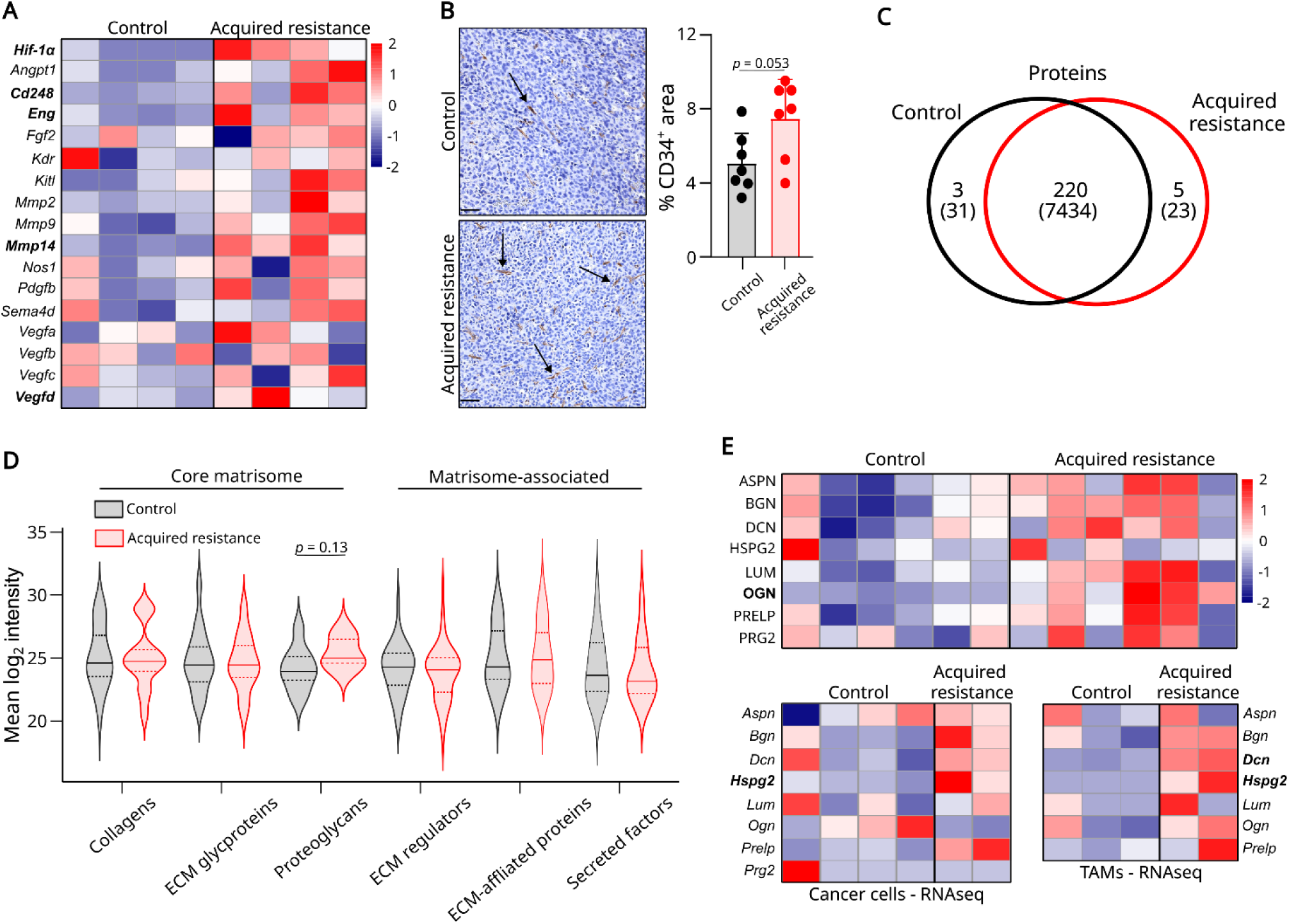
Acquired resistance involves hypoxia-induced angiogenesis and proteoglycan-enriched ECM remodeling. (A) Heatmap of normalized RNA-seq read counts for hypoxia-induced pro-angiogenic genes, scaled to Z-scores. (B) Representative CD34 immunohistochemistry in tissue from control and acquired resistance tumors, with a dot plot quantifying the DAB+ (brown) area. Scale bar: 50 µm. (C) Venn diagram illustrating the number of shared and distinct matrisome and total proteins between control and acquired resistance tumors (n = 6 per group). (D) Mean log₂ intensity values of detected matrisome proteins shown as a distribution with lines indicating the first quartile, median, and third quartile (n = 6 per group). (E) Heatmap of normalized proteoglycan abundance at the protein and RNA levels; protein intensities and RNA-seq values are Z-score scaled. Statistical comparisons were performed using a Wald test (A, E bottom) the Mann-Whitney test (B, D), and permutation-based multiple testing procedure (E top) with Benjamini–Hochberg correction (A, E). Differentially expressed proteins and transcripts are shown in bold. Bars represent means and error bars represent SD.

Transcriptomic analyses of resistant tumors revealed enrichment of ECM remodeling pathways, prompting further investigation of matrix composition. The ECM is a dynamic regulator of tumor architecture and immune cell function, and collagen deposition and matrix stiffening can limit T cell infiltration and directly suppress effector activity (28–30). Although ECM remodeling is implicated in tumor progression and primary resistance to immunotherapy (31), its role in acquired resistance remains poorly defined. Collagen abundance assessed by Picrosirius red staining did not differ between control and acquired resistance tumors (Supplementary Figure 4A), suggesting that collagen deposition alone does not account for ECM alterations in resistant tumors. To obtain a broader view of the ECM composition, matrisome-enriched tumor fragments were analyzed by mass spectrometry. A total of 7488 proteins were quantified, of which 228 were annotated as matrisome proteins (Figure 6C). Functional categorization revealed a trend toward increased abundance of proteoglycans in acquired resistance tumors (Figure 6D). Eight proteoglycans were identified, and their relative enrichment in resistant tumors was supported by proteomic and RNA-seq data (Figure 6E). As HIF-1α signaling has been linked to regulation of proteoglycan expression under hypoxic conditions (32,33), the observed accumulation of proteoglycans may reflect hypoxia-associated ECM remodeling in acquired resistance.

In addition to proteoglycans, several other ECM components exhibited altered expression profiles (Supplementary Figure 4B-D). Lysyl oxidase (LOX), an enzyme that cross-links collagen and elastin, was detected exclusively in acquired resistance tumors (Supplementary Figure 4D). Elastin (ELN), a substrate of LOX, was similarly present only in resistant tumors. In contrast, TIMP3 expression was absent in acquired resistance tumors (Supplementary Figure 4D). TIMP3 inhibits matrix metalloproteinases, and loss of TIMP3 has been linked to poor survival outcomes by permitting unchecked protease activity, contributing to ECM degradation (34). In combination with the RNA-seq analysis, which revealed elevated expression of the collagenolytic enzymes MMP2, MMP9, and MMP14 (Figure 6A), these data indicate enhanced matrix remodeling activity in acquired resistance tumors.

## DISCUSSION

Recently, patients with acquired resistance to ICIs have been recognized as a distinct patient group with their own distinct biological and clinical features, highlighting the need for novel therapeutic strategies to overcome resistance (3). This study presents a comprehensive *in vivo* model of acquired resistance to ICIs that recapitulates dynamic patterns of ICI therapeutic escape. We show that ICI-induced immune pressure drives metabolic reprogramming of cancer cells, leading to increased mitochondrial respiration. This hypermetabolic state promotes hypoxia in tumors with acquired resistance. Furthermore, ECM remodeling programs are induced, reshaping the tumor microenvironment. Although T cells infiltrate these tumors, the metabolically and structurally altered microenvironment promotes T cell dysfunction and facilitates tumor escape.

Unlike previous preclinical models that rely on repeated selection cycles of resistant cancer cells, our approach takes advantages of the intrinsic heterogeneous responses towards ICI treatment in MC38-tumor bearing mice, ranging from complete tumor regression to acquired and primary resistance. This heterogeneity resembles clinical responses and the complexity of secondary immune responses associated with ICI efficacy (3). Building on this observation, we explored whether the *in vivo* heterogeneity could serve as a functional proxy for clinical response patterns. To our knowledge, murine models rarely stratify individual mice based on differential responses to ICI therapy. Complete responders exhibited tumor clearance and durable immune memory. Tumors with acquired resistance did not show impaired T cell infiltration, a hallmark of primary resistance (35), suggesting distinct mechanisms. Our initial objective was to characterize the phenotype of acquired resistance. However, the stratifiable response groups suggest a broader potential: by treating mice individually rather than as experimental replicates, it may be possible to refine future models and address more complex questions, including why mice with primary resistance fail to generate an initial T cell response compared to mice with acquired resistance. While our study was not powered to resolve these questions due to cohort size, it lays the groundwork for such investigations.

Acquired genetic alterations have been implicated in individual patient cases of acquired resistance to ICIs (8–10,36). However, no recurrent or mechanistically relevant mutations associated with resistance were identified in our model. The generation of acquired resistance independently of genetic mutations is highly relevant, as a substantial proportion of patients treated with ICIs lack an identifiable genomic driver of resistance (11–13). Our model system could reveal mechanisms responsible for acquired resistance in some of these patient cases. The absence of genetic drivers of acquired resistance has also been reported in previous studies using murine models (37,38).

A broader characterization of the TME in acquired resistance showed that resistant tumors had a notable enrichment of TAMs. This accumulation of TAMs is consistent with previous studies reporting elevated TAM abundance in resistant states (16,39). The increase in TAMs is likely driven by enhanced recruitment to the TME, since cancer cells showed increased mRNA levels of chemokines such as CCL2. Transcriptomic profiling of cancer cells and TAMs in acquired resistance tumors revealed coordinated upregulation of pathways related to hypoxia, TNF-α signaling, and ECM remodeling. Hypoxia in tumors can arise through multiple mechanisms, including increased proliferative demand and metabolic activity (40). In line with this, we observed elevated basal oxygen consumption in cancer cells from acquired resistance tumors, suggesting enhanced mitochondrial respiration. Such a shift in metabolic phenotype likely contributes to immune dysfunction by reshaping the tumor microenvironment through nutrient competition, acidosis, and immunosuppressive signaling (41). While a previous syngeneic mouse model has linked metabolic changes to acquired resistance (15), this model relied on serial passaging to enforce resistance. In contrast, our model demonstrates that metabolic rewiring can emerge under ICI-induced immune pressure, without iterative selection. These findings extend the role of hypoxia in promoting immune evasion and primary resistance by showing that it can also emerge rapidly as an adaptive response to ICI-induced immune pressure. Recent studies have indeed begun to explore the role of hypoxia in the context of acquired resistance (38,42). Together, these findings strengthen the link between metabolic adaptation, hypoxia, and immune dysfunction in acquired resistance.

Transcriptomic analyses indicated changes in ECM-related pathways in resistant tumors, prompting a focused characterization of the ECM landscape. While the ECM has been increasingly implicated in immune modulation (29,43) and in primary resistance to ICI (31), its contribution to acquired resistance remains underexplored. Proteomic profiling revealed a trend towards increased levels of proteoglycans, including osteoglycin. These proteoglycans have been identified as mediators of tumor inflammation and immune suppression. Emerging evidence suggests that specific proteoglycan signatures influence response to immunotherapy in patients with colorectal cancer (43–45). Additionally, selective upregulation of LOX and elastin indicates active collagen crosslinking and matrix stiffening (46,47), structural alterations known to constrain immune cell motility and attenuate effector function. Together, these findings suggest that ICI-induced immune pressure promotes a mechanically and immunologically restrictive ECM state that supports tumor persistence. Recent studies, including work on COL6A1 and COL3A1 in the context of acquired resistance to anti-PD-1 therapy (48), support the notion that specific ECM components can influence immunotherapeutic outcomes and may serve as actionable biomarkers or therapeutic targets in future studies. Additional studies are required to determine whether ECM changes are causative or merely correlative in acquired resistance. Finally, complementary spatial and biomechanical analyses will be required to fully elucidate how ECM architecture regulates immune cell function *in vivo*.

Together, these data support a model in which ICIs initially suppress tumor growth, but sustained immunologic pressure promotes metabolic rewiring and hypoxia, which in turn promote ECM remodeling and immune dysfunction. This remodeled microenvironment creates a structurally and metabolically hostile niche for T cells, allowing tumor escape despite persistent infiltration. These findings have important implications for therapy. Proteoglycans and ECM regulators may serve as biomarkers for acquired resistance or as actionable targets. Furthermore, therapeutic strategies aimed at disrupting cancer cell metabolism and hypoxia, modulating ECM composition, or restoring immune cell function within remodeled niches may prove synergistic with ICIs. Future studies should investigate rational combination strategies to delay or overcome resistance, including the timing and sequencing of ICI and targeted agents.

## METHODS

### Cell lines

The mouse colon cancer cell line MC38 (Kerafast) was lentivirally transduced to express eGFP under control of the CMV promoter and cultured in Dulbecco’s Modified Eagle Medium (DMEM) supplemented with 10% fetal bovine serum (FBS), 1% penicillin-streptomycin (P/S), and 1 μg/mL puromycin (all from Gibco) under standard conditions. Additional cancer cell lines were established from control and acquired resistance tumors following enzymatic dissociation and *in vitro* expansion of maximally 8 passages. Cellular morphology and GFP expression were examined to confirm the identity as cancer cells.

### Murine tumor models

Animal experiments were conducted at the animal facility of the Department of Oncology, Copenhagen University Hospital - Herlev and Gentofte, under standard housing conditions. Female C57BL/6 (Taconic), B6.129S6-Rag2tm1FwaN12 (RAG2 KO, Taconic), and C57BL/6-Tg(TcraTcrb)1100Mjb/Crl (OT-I, Charles River) mice aged 8-20 weeks were used. Experimental procedures were approved by the Danish Animal Inspectorate. MC38 cells (5 × 10^5^) were injected subcutaneously into the flank. Tumor dimensions were measured three times weekly using a digital caliper and calculated as width^2^ x length x (π/6). Mice were euthanized when tumor volumes ≥1,200 mm^3^. Tumors and spleens were harvested for analysis. Tumor growth was analyzed using TumGrowth with default settings, which applies type II ANOVA and pairwise comparisons (49).

### Therapeutic treatments

When tumor volumes exceeded 50 mm^3^, mice were randomized 1:1 to control or treatment with anti-PD-1 monoclonal antibody (200 μg/mouse; clone RMP1-14, BioXCell) and anti-CTLA-4 monoclonal antibody (200 μg/mouse; clone 9D9, BioXCell) diluted in 100 uL PBS. Control-treated mice received PBS. Treatments were administered intraperitoneally three times during the first week, followed by weekly injections. Responses were defined based on survival. Acquired resistance was operationally defined as an initial delay in tumor growth followed by progression, resulting in survival exceeding 50% of the average survival of control-treated mice. This threshold was used to distinguish acquired resistance from primary resistance, as no standardized criteria exist for preclinical models, and serves as a proxy for treatment durability analogous to time-based clinical definitions (50).

### Retransplantation models

For retransplantation using cancer cells, tumor tissues were enzymatically digested and stained as described in Supplementary Methods. Cancer cells were sorted based on CD45^-^GFP^+^ expression on a BD FACSMelody™ cell sorter (BD Biosciences) and seeded into complete medium and maintained at 37°C in 5% CO₂ until reaching 80% confluency. Cancer cells underwent purity verification by flow cytometry before a total of 5 × 10⁵ cancer cells were injected subcutaneously into the flank of six recipient mice per tumor. For retransplantation using tumor fragments, recipient mice received ibuprofen (30 mg/kg; Reckitt) before surgery. Tumors were excised and sectioned into fragments (approx. 2 x 2 mm). Two fragments were transplanted subcutaneously into the right flank of each recipient via one skin incision. Postoperative pain management included daily ibuprofen for three days. Three tumors per group were transplanted into six recipients per tumor. Recipients were randomized 1:1 to control or ICI treatment.

### Enzyme-linked immune-spot (ELISpot) assay

Splenocytes from control- and ICI-treated mice were isolated by mechanical dissociation through a 70 μm cell strainer. Following RBC lysis for 5 minutes, cells were cryopreserved. An IFN-γ ELISPOT assay was performed using 96-well plates coated overnight at 4°C with an anti-mouse IFN-γ capture antibody (12 μg/mL; 3321-3-1000, MabTech). Plates were blocked with RPMI-1640 supplemented with 10% FBS and 1% P/S for 30 minutes at 37°C. A total of 8 × 10⁵ splenocytes/well were stimulated with the p15E peptide KSPWFTTL (5 μM; Schäfer-N). A positive control included stimulation with concanavalin A (5 μg/mL; C5275, Sigma Aldrich), while unstimulated cells were a negative control. After overnight incubation at 37°C, plates were developed using a biotinylated anti-mouse IFN-γ detection antibody (3321-6-250, MabTech), streptavidin-ALP (3310-10, MabTech), and BCIP/NBT substrate (3650-10, MabTech). Spots were quantified using a CTL ImmunoSpot S6 analyzer with ImmunoSpot software (v5.1, CTL Analyzers). Specific responses were calculated by subtracting unstimulated spot counts from peptide-stimulated values, with all conditions run in triplicate.

### Whole-genome sequencing

Genomic DNA was isolated from cancer cell lines using the AllPrep DNA/RNA Mini Kit (80204, Qiagen) according to the manufacturer’s protocol. Library preparation and sequencing were performed by the Genomic Medicine facility at Copenhagen University Hospital - Rigshospitalet. Whole-genome libraries were prepared using Illumina DNA PCRfree prep (20041795, Illumina) with an input of >300ng, followed by 150 bp paired-end sequencing on NovaSeq6000 platform, targeting a mean coverage depth of 30X. Raw reads were aligned to GRCm38 mouse reference genome using BWA-MEM, following the Sarek workflow (51,52). Single nucleotide variants and small insertions/deletions (indels) were identified with Strelka (53). Variant annotation was performed using Variant Effect Predictor (VEP), focusing on predicted deleterious mutations, including missense and nonsense mutations, splice site alterations, and frameshift indels. All possible 9-mer mutated peptides generated from nonsynonymous missense mutations were used to predict binding affinities to the C57BL/6 MHC class I alleles H-2K^b^ and H-2D^b^ with NetMHC (v4.0, DTU Health Tech) (54).

### Chromium-51 cytotoxicity assay

Cancer cell lines were pulsed with the MHC class I-restricted chicken ovalbumin-derived peptide SIINFEKL (1 μM; Schäfer-N) and labeled with 100 µCi of ⁵¹Cr for 1 hour at 37°C. SIINFEKL-specific CD8⁺ T were isolated from the spleen of an OT-1 mouse by mechanical dissociation through a 70 μm cell strainer and anti-CD8 magnetic bead separation (130-117-044, Miltenyi). Mouse T cells were expanded *in vitro* on plates pre-coated with anti-CD3 and anti-CD28 (5 mg/mL; BE0001-1 and BE0015-1, BioXCell) antibodies in complete RPMI-1640 with 1% ITS supplemented with IL-2 (20 ng/mL; 212-12, Peprotech), IL-7 (5 ng/mL; 577802, Biolegend), and IL-21 (10 ng/mL; 212-21, Peprotech) over four days. T cells and cancer cells were co-cultured for 4 hours at 37°C. Supernatants were collected, and the amount of ⁵¹Cr released was measured using a 2470 Automatic γ-counter (PerkinElmer). Maximum release was determined by lysing cancer cells with 10% Triton X-100, while spontaneous release was measured from cancer cells alone. Specific lysis was calculated by subtracting spontaneous release from experimental release and normalizing to the total releasable ⁵¹Cr (maximum minus spontaneous release). Assays were performed in technical triplicates, and maximum and spontaneous release wells with six technical replicates.

### Assessment of IFNγ responsiveness

Cancer cell lines were stimulated *in vitro* with IFNγ (20 ng/mL; 315-05, PeproTech) for 48 hours. Cells were harvested and stained for surface markers (antibody panels and staining conditions in Supplementary Methods). Flow cytometry was performed on a NovoCyte Quanteon flow cytometer (Agilent), and data were analyzed using NovoExpress software (v1.5, Agilent). Surface marker expression was reported as median fluorescence intensity (MFI) for PD-L1 and H-2Db, and as the percentage of live cells expressing I-A/I-E.

### RNA sequencing

RNA isolation was performed on sorted cancer cells, tumor-associated macrophages, and whole tumors. RNA samples were then subjected to poly-dT-enriched RNA sequencing on a NovaSeq 6000 system platform (Illumina). Differential gene expression analyses were performed in R (v4.3.0) using DESeq2 (adjusted p-value < 0.05; log₂ fold change > 0.585). Gene ontology and pathway enrichment analyses were performed using clusterProfiler. RNA-seq data are deposited in GEO (accession GSE325498). Detailed sample and library preparation, sequencing, and bioinformatic processing steps are described in the Supplementary Methods.

### Real-time *in vitro* proliferation

Cancer cell lines were seeded at a density of 10.000 cells/well in complete medium supplemented with 1.25 μg/mL fungizone in an E-Plate 96 (300601020, Agilent) and loaded onto the RTCA SP instrument (Agilent). Cell proliferation was monitored continuously for two days with impedance measurements recorded every 30 minutes. Data acquisition and analysis were performed using RTCA PRO software (v2.1.0, ACEA Biosciences). Assays were performed in technical triplicates, and proliferation rates were determined based on the dynamic changes in the Cell Index over time.

### Metabolic profiling

Cancer cell lines were seeded at a density of 50.000 cells/well in complete medium in an XF96 plate (101085-004, Agilent) and incubated overnight at 37°C. Confluency was confirmed, and the medium was replaced with Seahorse XF RPMI assay media (103576-100, Agilent) supplemented with 1 mM sodium pyruvate, 3 mM glutamine, and 4 mM glucose. Plates were equilibrated before being transferred to the XF PRO Analyzer (Agilent). Oxygen consumption rates (OCR) were recorded prior to sequential injection of mitochondrial stress test compounds: 2 µM oligomycin, 2 µM carbonyl cyanide p-(trifluoromethoxy) phenylhydrazone (FCCP), and 0.5 µM antimycin A. Assays were performed in 5-6 technical replicates. Data acquisition and analysis were performed using Seahorse Wave software (v2.6.3.8, Agilent) with OCR reported in pmol/min. ATP-linked respiration was calculated as the decrease in OCR after oligomycin addition, representing basal respiration coupled to ATP production. Maximal respiration was determined as the OCR after FCCP injection, while spare respiratory capacity was calculated as the difference between maximal and basal OCR.

### Phenotyping by flow cytometry

Single-cell suspensions were prepared from tumor fragments and stained for immune and stromal markers as detailed in the Supplementary Methods. Flow cytometry was performed on a NovoCyte Quanteon flow cytometer (Agilent) or FACS CantoII flow cytometer (BD Biosciences), and data were analyzed using NovoExpress software (v1.5, Agilent). Fluorescence minus one (FMO) controls were included where appropriate. All panels underwent a compensation procedure prior to analysis. Surface marker expression was reported as the percentage of live cells or the percentage of parent population.

### Acquired resistance signature

Genes driving hypoxia and EMT pathways in cancer cells and TAMs were identified by intersecting the leading-edge gene lists extracted from the GSEA output using the *core_enrichment* field. The overlapping genes were defined as the acquired resistance signature. Signature scores were analyzed in the TCGA colon adenocarcinoma (TCGA-COAD) dataset. For each sample, scores were calculated as the mean log₂-transformed expression of the signature genes. Resulting values were z-score normalized across samples to enable comparison between signatures and conditions. Correlations were computed between the signature z-scores and CD8⁺ T cell exhaustion scores obtained from the TIDE framework and associations with TIDE responder versus non-responder status were assessed (26).

### Immunohistochemistry

Tumors were fixed in 4% paraformaldehyde, embedded in paraffin, and cut into 5 μM sections according to standard methods. Hematoxylin and eosin (H&E) staining was performed according to standard protocols. For CD34 immunostaining, 10 mM citrate buffer, pH 6.0 antigen retrieval, and rat anti-mouse CD34 (1:400, MEC 14.7, Novus Biologicals) were used. The Dako mouse EnVision Detection system (Agilent) was used for detecting the primary antibody. For the detection of fibrillar collagen, sections were stained with 0.1% Sirius red (PSR) diluted in saturated picric acid (Ampliqon) and counterstained with hematoxylin. Images of PSR stained sections were acquired using a light microscope with polarization filters. The percentage of positive areas were quantified with QuPath software (v0.4.3). Positive and negative areas were manually assigned on several sections, and these were used to train a pixel classifier until it could reliably detect positive areas. The trained pixel classifier was run on the entire section (CD34) or ten squares randomly distributed across each section, blinded to the staining (PSR).

### ECM proteomics

Tumor tissues were subjected to ECM enrichment using the subcellular protein fractionation kit for tissues (87790, Thermo Scientific). The insoluble ECM fraction was processed for data-independent acquisition mass spectrometry on a Tribrid Ascend mass spectrometer (Thermo Scientific). Raw data were analyzed using DIA-NN (v.1.8.2) with a library predicted from Mus musculus Uniprot file (UP000000589) using a 1% false discovery rate (FDR) filtering. ECM proteins were annotated using the MatrisomeAnalyzeR tool (v1.0.1). Log₂-transformed protein intensities were used for differential expression analysis with permutation-based FDR correction (FDR < 0.05; log₂ fold change > 1). Detailed protein extraction, mass spectrometry acquisition parameters, and data processing workflows are provided in the Supplementary Methods.

## ACKNOWLEDGEMENTS

We would like to kindly thank the animal caretakers Anne Thorbek Nielsen and Ditte Stina Jensen at the Department of Oncology, Copenhagen University Hospital - Herlev and Gentofte, for their assistance related to the mouse studies. Whole genome sequencing was conducted at the MDxCore facility at Copenhagen University Hospital - Rigshospitalet. The analysis of whole genome sequencing data using the Sarek workflow was performed by the BRIC Bioinformatics Core Facility at the University of Copenhagen. Mass spectrometry based proteomic analysis and data analysis were performed by the Proteomics Research Infrastructure at the University of Copenhagen, supported by the Novo Nordisk Foundation (grant agreement number NNF19SA0059305).

## REFERENCES

1. Hellmann MD, Paz-Ares L, Bernabe Caro R, Zurawski B, Kim SW, Carcereny Costa E, et al. Nivolumab plus ipilimumab in advanced non–small-cell lung cancer. N Engl J Med. 2019 Nov 21;381(21):2020–31. doi:10.1056/NEJMoa1910231

2. Wolchok JD, Chiarion-Sileni V, Gonzalez R, Rutkowski P, Grob JJ, Cowey CL, et al. Overall survival with combined nivolumab and ipilimumab in advanced melanoma. N Engl J Med. 2017 Oct 5;377(14):1345–56. doi:10.1056/NEJMoa1709684

3. Schoenfeld AJ, Hellmann MD. Acquired resistance to immune checkpoint inhibitors. Cancer Cell. 2020 Apr 13;37(4):443–55. doi:10.1016/J.CCELL.2020.03.017

4. Zhuo N, Liu C, Zhang Q, Li J, Zhang X, Gong J, et al. Characteristics and prognosis of acquired resistance to immune checkpoint inhibitors in gastrointestinal cancer. JAMA Netw Open. 2022 Mar 1;5(3):e224637–e224637. doi:10.1001/JAMANETWORKOPEN.2022.4637

5. Liu F, Yin G, Tao Y, Pan Y. The efficacy of ICIs rechallenge in advanced small cell lung cancer after progression from ICIs plus chemotherapy: A real-world study. Int Immunopharmacol. 2025 Apr 16;152:114372. doi:10.1016/J.INTIMP.2025.114372

6. Wang B, Han Y, Zhang Y, Zhao Q, Wang H, Wei J, et al. Overcoming acquired resistance to cancer immune checkpoint therapy: potential strategies based on molecular mechanisms. Cell Biosci. 2023 Dec 1;13(1):120. doi:10.1186/s13578-023-01073-9

7. Marine JC, Dawson SJ, Dawson MA. Non-genetic mechanisms of therapeutic resistance in cancer. Nat Rev Cancer. 2020 Oct 8;20(12):743–56. doi:10.1038/S41568-020-00302-4

8. Gettinger S, Choi J, Hastings K, Truini A, Datar I, Sowell R, et al. Impaired HLA class I antigen processing and presentation as a mechanism of acquired resistance to immune checkpoint inhibitors in lung cancer. Cancer Discov. 2017 Dec 1;7(12):1420–35. doi:10.1158/2159-8290.CD-17-0593

9. Nielsen M, Presti M, Sztupinszki Z, Jensen AWP, Draghi A, Chamberlain CA, et al. Co-existing alterations of MHC class I antigen presentation and IFNγ signaling mediate acquired resistance of melanoma to post-PD-1 immunotherapy. Cancer Immunol Res. 2022 Oct 4;10(10):1254–62. doi:10.1158/2326-6066.CIR-22-0326

10. Zaretsky JM, Garcia-Diaz A, Shin DS, Escuin-Ordinas H, Hugo W, Hu-Lieskovan S, et al. Mutations associated with acquired resistance to PD-1 blockade in melanoma. N Engl J Med. 2016 Sep 1;375(9):819–29. doi:10.1056/NEJMOA1604958

11. Hiltbrunner S, Cords L, Kasser S, Freiberger SN, Kreutzer S, Toussaint NC, et al. Acquired resistance to anti-PD1 therapy in patients with NSCLC associates with immunosuppressive T cell phenotype. Nat Commun. 2023 Aug 24;14(1):5154. doi:10.1038/S41467-023-40745-5

12. Memon D, Schoenfeld AJ, Ye D, Fromm G, Rizvi H, Zhang X, et al. Clinical and molecular features of acquired resistance to immunotherapy in non-small cell lung cancer. Cancer Cell. 2024 Feb 12;(42):1–16. doi:10.1016/j.ccell.2023.12.013

13. Schiantarelli J, Benamar M, Park J, Sax HE, Oliveira G, Bosma-Moody A, et al. Genomic mediators of acquired resistance to immunotherapy in metastatic melanoma. Cancer Cell. 2025 Feb 10;43(2):308–16. doi:10.1016/J.CCELL.2025.01.009

14. Verdys P, Johansen AZ, Gupta A, Presti M, Dionisio E, Madsen DH, et al. Acquired resistance to immunotherapy in solid tumors. Trends Mol Med. 2025 Nov;11(31):1008–20. doi:10.1016/J.MOLMED.2025.03.010

15. Jaiswal AR, Liu AJ, Pudakalakatti S, Dutta P, Jayaprakash P, Bartkowiak T, et al. Melanoma evolves complete immunotherapy resistance through the acquisition of a hypermetabolic phenotype. Cancer Immunol Res. 2020 Nov 1;8(11):1365. doi:10.1158/2326-6066.CIR-19-0005

16. Denis M, Grasselly C, Choffour PA, Wierinckx A, Mathé D, Chettab K, et al. In vivo syngeneic tumor models with acquired tesistance to anti-PD-1/PD-L1 therapies. Cancer Immunol Res. 2022 Aug 1;10(8):1013–27. doi:10.1158/2326-6066.CIR-21-0802

17. Koyama S, Akbay EA, Li YY, Herter-Sprie GS, Buczkowski KA, Richards WG, et al. Adaptive resistance to therapeutic PD-1 blockade is associated with upregulation of alternative immune checkpoints. Nat Commun. 2016 Feb 17;7(1):1–9. doi:10.1038/ncomms10501

18. Zandberg DP, Menk A V., Velez M, Normolle D, Depeaux K, Liu A, et al. Tumor hypoxia is associated with resistance to PD-1 blockade in squamous cell carcinoma of the head and neck. J Immunother Cancer. 2021 May 1;9(5):e002088. doi:10.1136/JITC-2020-002088

19. Bernardo M, Tolstykh T, Zhang Y an, Bangari DS, Cao H, Heyl KA, et al. An experimental model of anti-PD-1 resistance exhibits activation of TGFß and Notch pathways and is sensitive to local mRNA immunotherapy. Oncoimmunology. 2021;10(1):1881268. doi:10.1080/2162402X.2021.1881268

20. Xiong Z, Chan SL, Zhou J, Vong JSL, Kwong TT, Zeng X, et al. Targeting PPAR-gamma counteracts tumour adaptation to immune-checkpoint blockade in hepatocellular carcinoma. Gut. 2023 Sep 1;72(9):1758–73. doi:10.1136/gutjnl-2022-328364

21. Efremova M, Rieder D, Klepsch V, Charoentong P, Finotello F, Hackl H, et al. Targeting immune checkpoints potentiates immunoediting and changes the dynamics of tumor evolution. Nat Commun. 2018 Jan 2;9(1):1–13. doi:10.1038/s41467-017-02424-0

22. Wei SC, Anang NAAS, Sharma R, Andrews MC, Reuben A, Levine JH, et al. Combination anti–CTLA-4 plus anti–PD-1 checkpoint blockade utilizes cellular mechanisms partially distinct from monotherapies. Proc Natl Acad Sci U S A. 2019 Nov 5;116(45):22699–709. doi:10.1073/pnas.1821218116

23. Zeh HJ, Perry-Lalley D, Dudley ME, Rosenberg SA, Yang JC. High avidity CTLs for two self-antigens demonstrate superior in vitro and in vivo antitumor efficacy. J Immunol. 1999 Jan 15;162(2):989–94. doi:10.4049/jimmunol.162.2.989

24. Zhong W, Myers JS, Wang F, Wang K, Lucas J, Rosfjord E, et al. Comparison of the molecular and cellular phenotypes of common mouse syngeneic models with human tumors. BMC Genomics. 2020 Jan 2;21(1):2. doi:10.1186/s12864-019-6344-3

25. Schrörs B, Hos BJ, Yildiz IG, Löwer M, Lang F, Holtsträter C, et al. MC38 colorectal tumor cell lines from two different sources display substantial differences in transcriptome, mutanome and neoantigen expression. Front Immunol. 2023 Mar 8;14(14):1102282. doi:10.3389/fimmu.2023.1102282

26. Jiang P, Gu S, Pan D, Fu J, Sahu A, Hu X, et al. Signatures of T cell dysfunction and exclusion predict cancer immunotherapy response. Nat Med. 2018 Oct 1;24(10):1550–8. doi:10.1038/S41591-018-0136-1

27. Krock BL, Skuli N, Simon MC. Hypoxia-induced angiogenesis: Good and evil. Genes Cancer. 2011 Dec;2(12):1117. doi:10.1177/1947601911423654

28. Kuczek DE, Larsen AMH, Thorseth ML, Carretta M, Kalvisa A, Siersbæk MS, et al. Collagen density regulates the activity of tumor-infiltrating T cells. J Immunother Cancer. 2019 Mar 12;7(1):68. doi:10.1186/s40425-019-0556-6

29. Rømer AMA, Thorseth ML, Madsen DH. Immune modulatory properties of collagen in cancer. Front Immunol. 2021 Dec 8;12:791453. doi:10.3389/fimmu.2021.791453

30. Salmon H, Franciszkiewicz K, Damotte D, Dieu-Nosjean MC, Validire P, Trautmann A, et al. Matrix architecture defines the preferential localization and migration of T cells into the stroma of human lung tumors. J Clin Invest. 2012 Mar 1;122(3):899–910. doi:10.1172/JCI45817

31. Hu Q, Zhu Y, Mei J, Liu Y, Zhou G. Extracellular matrix dynamics in tumor immunoregulation: from tumor microenvironment to immunotherapy. J Hematol Oncol. 2025 Dec 1;18(1):65. doi:10.1186/S13045-025-01717-Y

32. Asplund A, Stillemark-Billton P, Larsson E, Rydberg EK, Moses J, Hultén LM, et al. Hypoxic regulation of secreted proteoglycans in macrophages. Glycobiology. 2010;20(1):33–40. doi:10.1093/glycob/cwp139

33. Sotoodehnejadnematalahi F, Staples KJ, Chrysanthou E, Pearson H, Ziegler-Heitbrock L, Burke B. Mechanisms of hypoxic up-regulation of versican gene expression in macrophages. PLoS One. 2015 Jun 9;10(6):e0125799. doi:10.1371/journal.pone.0125799

34. Winer A, Adams S, Mignatti P. Matrix metalloproteinase inhibitors in cancer therapy: Turning past failures into future successes. Mol Cancer Ther. 2018 Jun 1;17(6):1147–55. doi:10.1158/1535-7163.MCT-17-0646

35. Sharma P, Hu-Lieskovan S, Wargo JA, Ribas A. Primary, adaptive, and acquired resistance to cancer immunotherapy. Cell. 2017 Feb 9;168(4):707–23. doi:10.1016/j.cell.2017.01.017

36. Sade-Feldman M, Jiao YJ, Chen JH, Rooney MS, Barzily-Rokni M, Eliane JP, et al. Resistance to checkpoint blockade therapy through inactivation of antigen presentation. Nat Commun. 2017 Oct 26;8(1):1–11. doi:10.1038/s41467-017-01062-w

37. Beyranvand Nejad E, Labrie C, Abdulrahman Z, Van Elsas MJ, Rademaker E, Kleinovink JW, et al. Lack of myeloid cell infiltration as an acquired resistance strategy to immunotherapy. J Immunother Cancer. 2020 Sep 1;8(2):e001326. doi:10.1136/JITC-2020-001326

38. Robles-Oteíza C, Hastings K, Choi J, Sirois I, Ravi A, Expósito F, et al. Hypoxia is linked to acquired resistance to immune checkpoint inhibitors in lung cancer. J Exp Med. 2025 Jan 6;222(1):e20231106. doi:10.1084/jem.20231106

39. Taylor MA, Hughes AM, Walton J, Coenen-Stass AML, Magiera L, Mooney L, et al. Longitudinal immune characterization of syngeneic tumor models to enable model selection for immune oncology drug discovery. J Immunother Cancer. 2019 Nov 28;7(1):328. doi:10.1186/S40425-019-0794-7

40. Chen Z, Han F, Du Y, Shi H, Zhou W. Hypoxic microenvironment in cancer: molecular mechanisms and therapeutic interventions. Signal Transduct Target Ther. 2023 Dec 1;8(1):70. doi:10.1038/S41392-023-01332-8

41. Li S, Zhang Y, Tong H, Sun H, Liao H, Li Q, et al. Metabolic regulation of immunity in the tumor microenvironment. Cell Rep. 2025 Nov 25;44(11):116463. doi:10.1016/j.celrep.2025.116463

42. Estephan H, Tailor A, Parker R, Kreamer M, Papandreou I, Campo L, et al. Hypoxia promotes tumor immune evasion by suppressing MHC-I expression and antigen presentation. EMBO Journal. 2025 Feb 3;44(3):903–22. doi:10.1038/S44318-024-00319-7

43. Sutherland TE, Dyer DP, Allen JE. The extracellular matrix and the immune system: A mutually dependent relationship. Science (1979). 2023 Feb 17;379:6633. doi:10.1126/science.abp8964

44. Deb G, Cicala A, Papadas A, Asimakopoulos F. Matrix proteoglycans in tumor inflammation and immunity. Am J Physiol Cell Physiol. 2022 Sep 1;323(3):C678–93. doi:10.1152/AJPCELL.00023.2022

45. Deming DA, Kraus SG, Brand J, Johnson KA, Abbott D, Kratz J, et al. Tumor matrix proteoglycan accumulation and processing alters T cell effector function and the response to immunotherapy in patients with oligometastatic colorectal cancer. Clin Cancer Res. 2026 Feb 11;Online ahead of print. doi:10.1158/1078-0432.CCR-25-2780

46. Joshi A, Dutta A, Roy A, Mondal B, Basak T. Decoding ECM remodeling: proteases as gatekeepers of Lysyl oxidase family-mediated crosslinking. FEBS Journal. 2026 Jan 28;Online ahead of print. doi:10.1111/febs.70418

47. Barker HE, Cox TR, Erler JT. The rationale for targeting the LOX family in cancer. Nat Rev Cancer. 2012 Aug;12(8):540–52. doi:10.1038/nrc3319

48. Wang M, Wang Y, Pan X, Wang B, Wang Y, Luo X, et al. Acquired resistance to immunotherapy by physical barriers with cancer cell–expressing collagens in non–small cell lung cancer. Proc Natl Acad Sci U S A. 2025 Jun 17;122(24):e2500019122. doi:10.1073/PNAS.2500019122

49. Enot DP, Vacchelli E, Jacquelot N, Zitvogel L, Kroemer G. TumGrowth: An open-access web tool for the statistical analysis of tumor growth curves. Oncoimmunology. 2018;7(9):e1462431. doi:10.1080/2162402X.2018.1462431

50. Kluger HM, Tawbi H, Feltquate D, LaVallee T, Rizvi NA, Sharon E, et al. Society for Immunotherapy of Cancer (SITC) checkpoint inhibitor resistance definitions: efforts to harmonize terminology and accelerate immuno-oncology drug development. J Immunother Cancer. 2023 Jul 1;11(7):e007309. doi:10.1136/JITC-2023-007309

51. Garcia M, Juhos S, Larsson M, Olason PI, Martin M, Eisfeldt J, et al. Sarek: A portable workflow for whole-genome sequencing analysis of germline and somatic variants. F1000Res. 2020 Sep 4;9:63. doi:10.12688/f1000research.16665.2

52. Hanssen F, Garcia MU, Folkersen L, Pedersen AS, Lescai F, Jodoin S, et al. Scalable and efficient DNA sequencing analysis on different compute infrastructures aiding variant discovery. bioRxiv. 2024 Feb 14;549462. doi:10.1101/2023.07.19.549462

53. Kim S, Scheffler K, Halpern AL, Bekritsky MA, Noh E, Källberg M, et al. Strelka2: fast and accurate calling of germline and somatic variants. Nat Methods. 2018 Jul 16;15(8):591–4. doi:10.1038/s41592-018-0051-x

54. Andreatta M, Nielsen M. Gapped sequence alignment using artificial neural networks: Application to the MHC class i system. Bioinformatics. 2016 Feb 15;32(4):511–7. doi:10.1093/BIOINFORMATICS/BTV639

