## Supplementary File for "Acquired resistance to immune checkpoint inhibitors is associated with hypoxia and ECM remodeling in colorectal cancer"

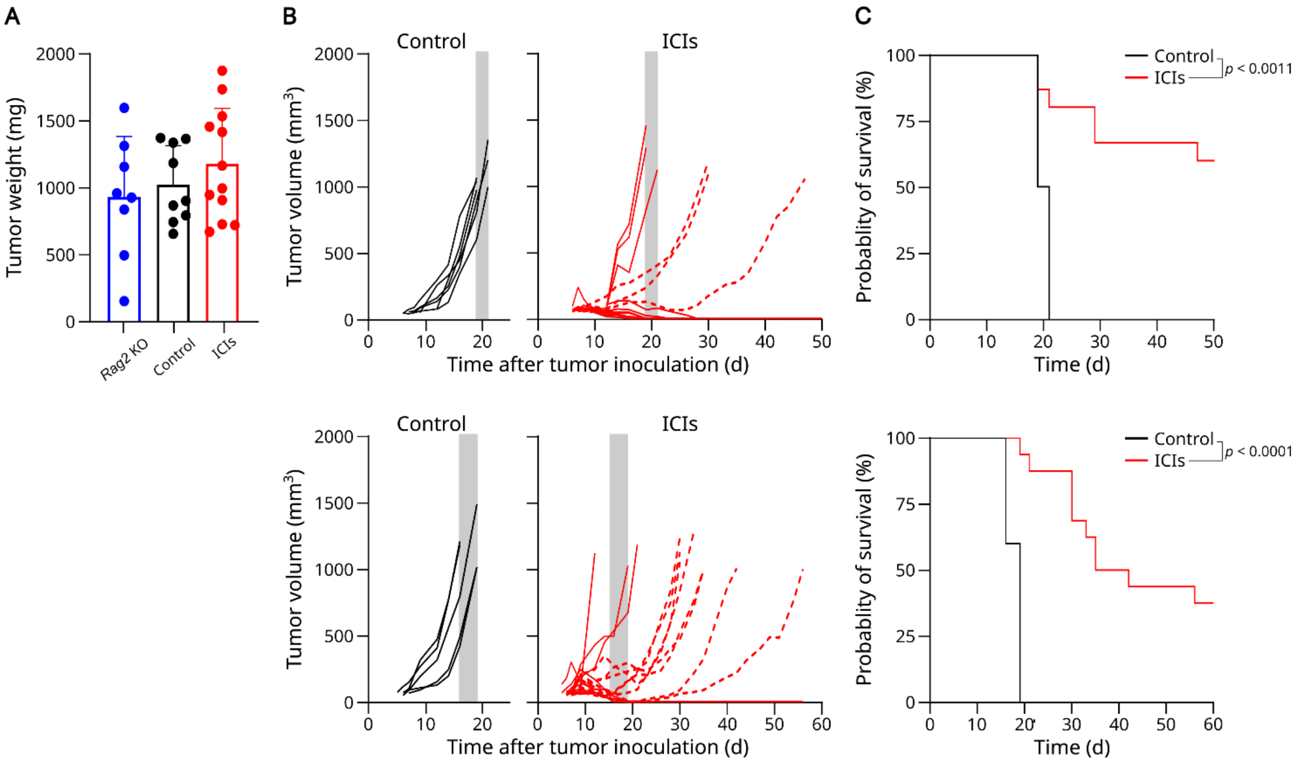


**Supplementary Figure 1. Comparable treatment outcomes across independent mouse experiments.** (A) Tumor weights at the study endpoint. (B) Individual tumor growth curves from two independent experiments. Grey box represents the terminal tumor volume time frame of the control group. Dotted lines indicate tumors that developed acquired resistance. (C) Survival curves for control-treated (n = 6 in the first experiment, n = 5 in the second) and ICI-treated wildtype C57BL/6 mice (n = 18 in both experiments). Differences in survival were assessed with the log-rank test. Bars represent means and error bars represent SD.

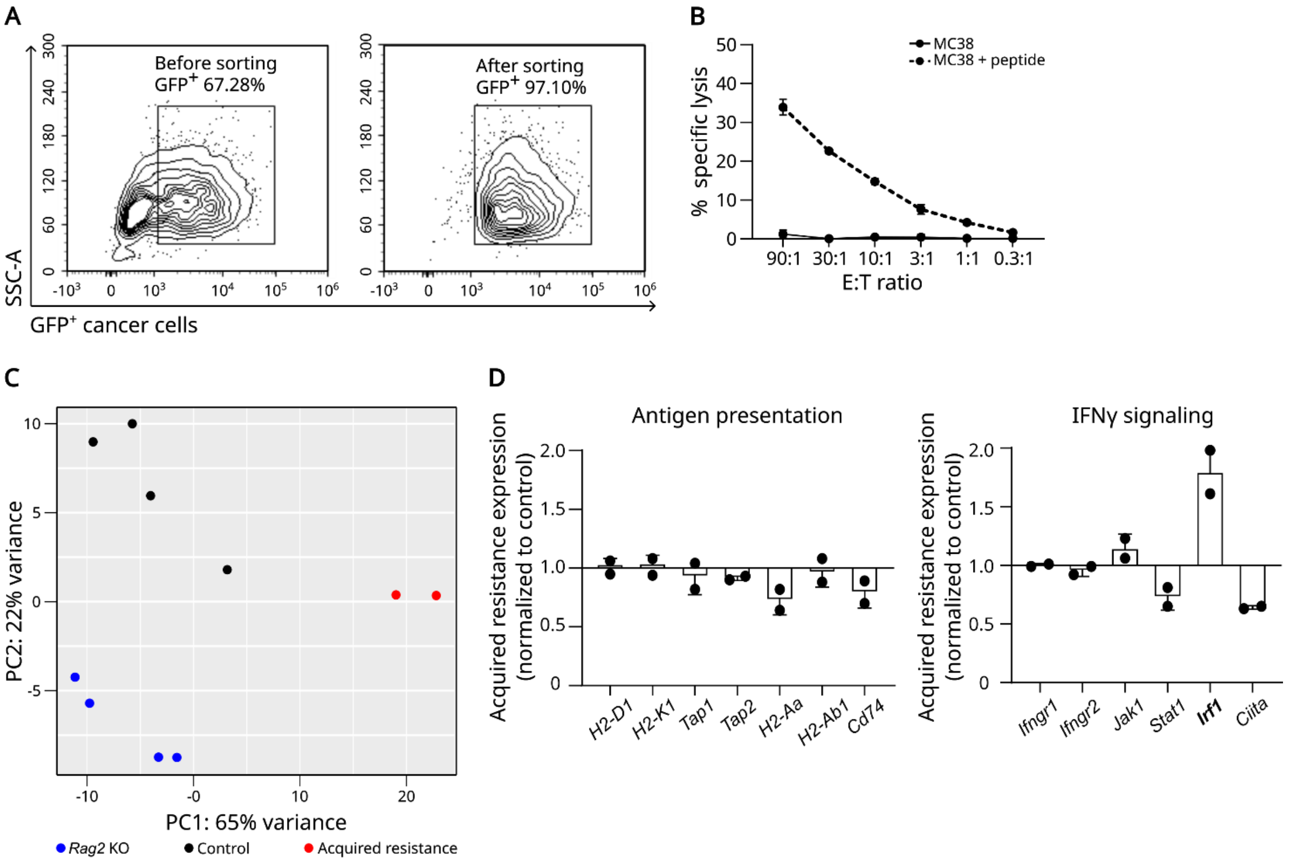


E

| HALLMARK ID | NES | Adjusted *p* value |
|---|---|---|
| TNFA SIGNALING VIA NFKB | 1.80 | 2.4e$^{-9}$ |
| ALLOGRAFT REJECTION | 1.74 | 7.6e$^{-8}$ |
| INFLAMMATORY RESPONSE | 1.74 | 7.6e$^{-8}$ |
| KRAS SIGNALING UP | 1.72 | 1.5e$^{-7}$ |
| EPITHELIAL MESENCHYMAL TRANSITION | 1.62 | 2.5e$^{-5}$ |
| COMPLEMENT | 1.55 | 4.1e$^{-4}$ |
| IL2 STAT5 SIGNALING | 1.49 | 2.8e$^{-3}$ |
| APOPTOSIS | 1.49 | 3.6e$^{-3}$ |
| HYPOXIA | 1.47 | 3.6e$^{-3}$ |
| COAGULATION | 1.48 | 4.7e$^{-3}$ |
| IL6 JAK STAT3 SIGNALING | 1.48 | 1.4e$^{-2}$ |
| UV RESPONSE UP | 1.41 | 1.4e$^{-2}$ |
| APICAL JUNCTION | 1.38 | 1.4e$^{-2}$ |
| INTERFERON GAMMA RESPONSE | 1.39 | 1.6e$^{-2}$ |
| P53 PATHWAY | 1.36 | 2.5e$^{-2}$ |
| KRAS SIGNALING DN | 1.35 | 4.8e$^{-2}$ |

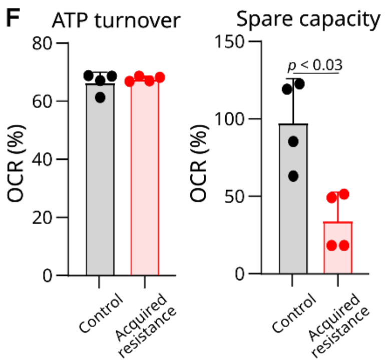


**Supplementary Figure 2. Cancer cells.** (A) Representative contour plots of GFP$^+$ cancer cells before and after sorting. (B) Percentages of the specific lysis of MC38 cell line ± SIINFEKL peptide. (C) PCA plot. (D) Expression of genes encoding antigen processing and presentation and IFNγ signaling in cancer cells, shown as RNA-seq counts normalized to average control expression. (E) Overview of significant hallmark gene sets. (F) Bar graphs of ATP turnover and spare respiratory capacity. Statistical comparisons were performed using a permutation-based multiple testing procedure with Benjamini-Hochberg correction (E) and the Mann-

Whitney test (F). Differentially expressed transcripts are shown in bold. Bars represent means and error bars represent SD.

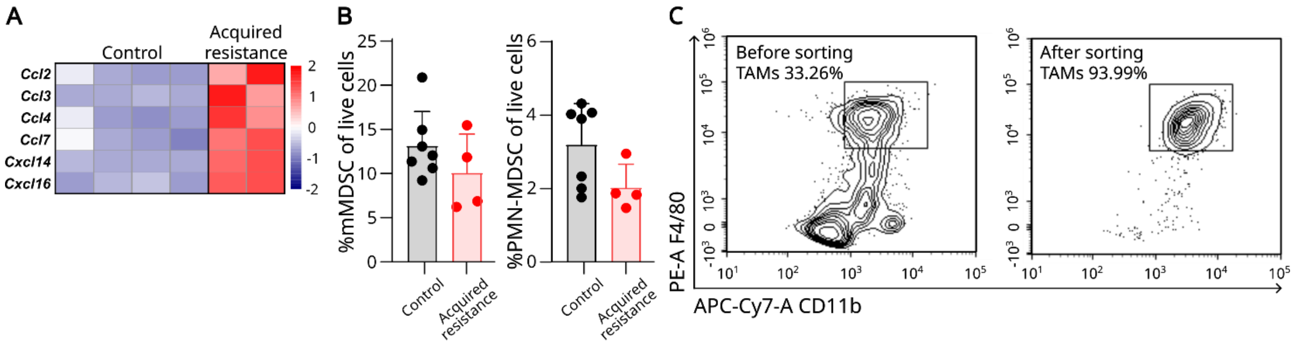


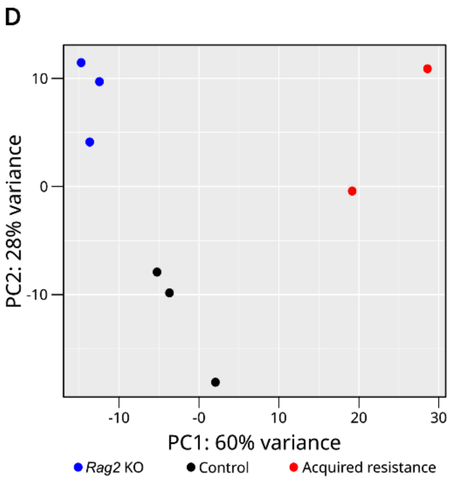


E

| HALLMARK ID | NES | Adjusted *p* value |
|---|---|---|
| EPITHELIAL MESENCHYMAL TRANSITION | 2.14 | $8.0e^{-11}$ |
| UV RESPONSE UP DOWN | 1.95 | $1.8e^{-6}$ |
| MYOGENESIS | 1.84 | $2.2e^{-6}$ |
| APICAL JUNCTION | 1.82 | $2.2e\text{-}^{6}$ |
| MYC TARGETS V1 | 1.76 | $2.2e^{-6}$ |
| EF2 TARGETS | 1.74 | $2.6e^{-6}$ |
| ESTROGEN RESPONSE EARLY | 1.78 | $4.4e^{-6}$ |
| ESTROGEN RESPONSE LATE | 1.77 | $1.0e^{-5}$ |
| ADIPOGENESIS | 1.68 | $2.6e^{-5}$ |
| P53 PATHWAY | 1.66 | $2.6e^{-5}$ |
| HYPOXIA | 1.64 | $6.1e^{-5}$ |
| TNFA SIGNALING VIA NFKB | 1.60 | $1.2e^{-4}$ |
| G2M CHECKPOINT | 1.58 | $1.5e^{-4}$ |
| CHOLESTEROL HOMEOSTASIS | 1.77 | $2.4e^{-4}$ |
| MTORC1 SIGNALING | 1.52 | $3.2e^{-4}$ |
| APOPTOSIS | 1.52 | $3.2e^{-4}$ |
| APICAL SURFACE | 1.88 | $8.4e^{-4}$ |
| KRAS SIGNALING UP | 1.51 | $8.8e^{-4}$ |
| PANCREAS BETA CLLS | 1.86 | $1.8e^{-3}$ |
| PEROXISOME | 1.62 | 1.9e-3 |
| MYC TARGETS V2 | 1.62 | $2.1e^{-3}$ |
| COAGULATION | 1.53 | $2.1e^{-3}$ |
| XENOBIOTIC METABOLISM | 1.42 | $3.1e^{-3}$ |
| FATTY ACID METABOLISM | 1.46 | $4.7e^{-3}$ |
| GLYCOLYSIS | 1.40 | $4.7e^{-3}$ |
| UV RESPONSE UP | 1.37 | $1.2e^{-2}$ |
| UNFOLDED PROTEIN RESPONSE | 1.40 | $1.2e^{-2}$ |
| TGF BETA SIGNALING | 1.55 | $1.8e^{-2}$ |
| IL2 STAT5 SIGNALING | 1.28 | $2.1e^{-2}$ |

**Supplementary Figure 3. Tumor immune microenvironment.** (A) Heatmap of normalized RNA-seq read counts for chemokines in cancer cells, scaled to Z-scores. (B) Bar graphs of mMDSCs ($CD45^+CD11b^+F4/80^-Ly6C^+Ly6G^-$) and PMN-MDSCs ($CD45^+$ $CD11b^+F4/80^-$ $Ly6C^{low}Ly6G^+$) in tumors harvested from control-treated and ICI-treated mice with acquired resistance. (C) Contour plots of TAMs ($CD45^+CD11b^+F4/80^+$) before and after sorting. (D) PCA plot. (E) Overview of significant hallmark gene sets. Statistical comparisons were performed using a Wald test (A) and a permutation-based multiple testing procedure with Benjamini-Hochberg correction (A, E). Differentially expressed transcripts are shown in bold. Bars represent means and error bars represent SD.

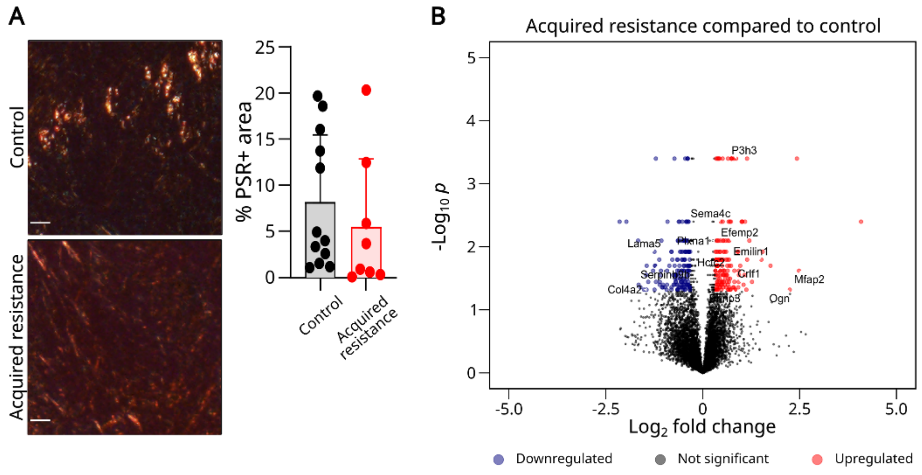


**C**

| Protein | Matrisome Category |
|---|---|
| **Down in acquired resistance** | |
| LAMA5 | ECM glycoproteins |
| SERPINB9B | ECM regulators |
| COL4A2 | Collagens |
| **Up in acquired resistance** | |
| EFEMP2 | ECM glycoproteins |
| EMILIN1 | ECM glycoproteins |
| MFAP2 | ECM glycoproteins |
| OGN | Proteoglycans |
| PLXNA1 | ECM-affiliated proteins |
| SEMA4C | ECM-affiliated proteins |
| P3H3 | ECM regulators |
| HCFC2 | Secreted factors |
| BRINP3 | Secreted factors |
| CRLF1 | Secreted factors |

**D**

| Protein | Matrisome Category |
|---|---|
| **Lost in acquired resistance** | |
| GDF15 | Secreted factors |
| MBL1 | ECM-affiliated proteins |
| TIMP3 | ECM regulators |
| **Gained in acquired resistance** | |
| ELN | ECM glycoproteins |
| MFAP5 | ECM glycoproteins |
| TNXB | ECM glycoprotein |
| ANXA8 | ECM-affiliated proteins |
| LOX | ECM regulators |

**Supplementary Figure 4. Extracellular matrix.** (A) Representative picrosirius red (PSR; red) staining in MC38 tumor tissue from control- and ICI-tumors with acquired resistance, with a dot plot showing quantification of PSR+ area. Scale bar: 50 µm. (B) Differentially expressed proteins (fold change > +/-1.25, *p* value < 0.05) in ICI-treated tumors with acquired resistance compared to control-treated (n = 6 per group). (C) Overview of differentially expressed matrisome proteins. (D) Overview of matrisome proteins lost (expression detected in ≤1/6 tumors with acquired resistance, ≥4/6 control tumors) or gained (expression detected in ≥4/6 tumors with acquired resistance, ≤1/6 control tumors) in ICI-treated tumors with acquired resistance. Statistical comparisons were performed using an unpaired t-test with permutation-based FDR correction (B-C). Bars represent means and error bars represent SD.

**Supplementary Table 1.** Known mutations and neoantigens in the MC38 cell line (source: Kerafast).

| **Mutations** | | | |
|---|---|---|---|
| **Gene** | **AA position** | **Type** | **MutPosition** |
| *Abl1* | K7N | Missense | 2:31689827 |
| *Arid1a* | Q602H | Missense & splice region | 4:133720550 |
| *Braf* | W487C | Missense | 6:39643182 |
| *Sox9* | T379R | Missense | 11:112785122 |
| *Trp53* | G242V | Missense | 11:69589202 |
| *Trp53* | S258I | Missense & splice region | 11:69589250 |
| **Neoantigens** | | | |
| **Gene** | **AA position** | **Mutated sequence** | **MutPosition** |
| *Aatf* | A500T | MAPIDHT**T**M | 11:8444 |
| *Cpne1* | D302Y | SSP**Y**SLHYL | 2:15607 |
| *Dpagt1* | - | - | - |
| *Tdg* | - | - | - |

- Indicates that the neoantigen was not found in this dataset.

**Supplementary Table 3.** Genes in acquired resistance signature.

| **Gene** | **Hallmark signature** | **Cancer cells** | | **TAMs** | |
|---|---|---|---|---|---|
| | | log2 FC | FDR | log2 FC | FDR |
| *Ackr3* | Hypoxia | 1.66 | 1.87E-06 | 5.67 | 2.03E-05 |
| *Bhlhe40* | Hypoxia | 2.17 | 1.00E-10 | 1.33 | 0.0008 |
| *Ddit4* | Hypoxia | 0.73 | 1.10E-08 | 1.69 | 0.0007 |
| *Errfi1* | Hypoxia | 1.28 | 0.0083 | 0.76 | 0.0196 |
| *Fbn1* | EMT | 0.82 | 0.0355 | 2.12 | 4.12E-09 |
| *Gem* | EMT | 2.41 | 1.12E-08 | 2.74 | 3.78E-07 |
| *Has1* | Hypoxia | 2.66 | 3.04E-07 | 1.88 | 0.0430 |
| *Hspa5* | Hypoxia | 0.84 | 1.06E-18 | 0.83 | 0.0005 |
| *Igfbp4* | EMT | 1.31 | 3.64E-11 | 2.85 | 1.30E-15 |
| *Lamc2* | EMT | 1.15 | 9.86E-07 | 4.02 | 7.33E-06 |
| *Loxl2* | EMT | 1.19 | 0.0006 | 2.68 | 0.0154 |
| *Mgp* | EMT | 0.93 | 0.0002 | 3.98 | 1.92E-08 |
| *Mylk* | EMT | 1.21 | 0.0059 | 3.51 | 3.10E-09 |
| *Rhob* | EMT | 2.42 | 0.0003 | 1.21 | 0.0011 |
| *Serpine1* | EMT and Hypoxia | 1.19 | 0.0027 | 2.29 | 1.18E-09 |
| *Srpx* | Hypoxia | 2.15 | 6.62E-05 | 7.99 | 0.0034 |

FC = fold change; FDR = false discovery rate.

**Supplementary Methods**

**Detailed flow cytometry**

*Retransplantation models*

For retransplantation using dissociated cancer cells, tumor tissues were chopped to small fragments in RPMI-1640 medium (Gibco) containing 1% P/S, collagenase type I (2.1 mg/mL; Gibco), and DNase I (75 µg/mL; Sigma Aldrich). The mixture was digested at 37°C for 45 minutes, filtered through a 70-µm cell strainer, treated with red blood cell (RBC) lysis buffer (Qiagen) for 5 minutes, and washed with RPMI-1640 medium supplemented with 10% FBS and 1% P/S. Single-cell suspensions were blocked with anti-FcγRIII/II antibody (Miltenyi) at 4°C for 10 minutes before surface staining in staining buffer with the following antibodies for 30 minutes at 4°C: CD45:Pe-Cy7 (1 µL/$10^6$ cells; 103114, Biolegend). Viability was assessed using Zombie Aqua Dye (77143, BioLegend).

*Assessment of IFN-γ responsiveness*

Harvested cancer cells were stained in staining buffer for 30 minutes at 4°C using the following antibodies: PD-L1:APC (1:50; 564715, BD Biosciences), H-2Db:APC-Fire (1:50; 114618, Biolegend), and I-A/I-E:PE-Cy7 (1:50; 107630, Biolegend). Cell viability was assessed using Zombie Aqua Dye (77143, BioLegend).

*Phenotyping by flow cytometry*

Single-cell suspensions were blocked with anti-FcγRIII/II antibody (Miltenyi) at 4°C for 10 minutes before surface staining in staining buffer for 20 minutes at 4°C using the following antibodies: CD3:FITC (17A2, 100204), CD4:BV421 (GK1.5, 100438), CD8:APC (53-6.7, 100712), CD11b: APC-Cy7 (M1/M70, 101226), CD44:APC-Cy7 (IM7, 103028), CD45:BV605 (30-F11, 103140), CD45:Pe-Cy7 (30-F11, 103114), CD62L:BV650 (MEL-1, 564108, BD Bioscience), F4/80:BB700 (T45-2342, 746070, BD Bioscience), biotinylated FAP (BAF3715, R&D Systems), Ly6C:AF700 (HK1.4, 128024), Ly6G:BV785 (1A8, 127645). All antibodies were from BioLegend and used 1:100 dilutions unless otherwise stated. For biotinylated antibodies, cells were stained with APC-conjugated streptavidin (405207, Biolegend) in staining buffer for 10 minutes at 4°C. Dead cells were excluded using Zombie Aqua Dye (77143, Biolegend).

**Detailed RNA sequencing**

*Sample preparation*

Single-cell suspensions from tumor tissues were prepared as previously described. Single-cell suspensions were blocked with anti-FcγRIII/II antibody (Miltenyi) at 4°C for 10 minutes before surface staining in staining buffer for 30 minutes at 4°C using the following antibodies: CD45:Pe-Cy7 (1.25 μL/$10^6$ cells; 103114, Biolegend), CD11b:APC-Cy7 (1.25 μL/$10^6$ cells; 101226, Biolegend), and F4/80:PE (1.25 μL/$10^6$ cells; 123110, Biolegend). Cell viability was assessed using Zombie Aqua Dye (77143, BioLegend). Cell sorting was performed on a BD FACSMelody™ cell sorter (BD Biosciences). Cancer cells were identified as $CD45^-GFP^+$, while macrophages were sorted as $CD45^+CD11b^+F4/80^+$. Purity was assessed immediately post-sorting. Sorted cells were stored at -80°C until RNA extraction using the RNeasy Mini Kit (74104, Qiagen) according to the manufacturer's protocol. For bulk tumor tissue RNA sequencing, tumor tissues were preserved in RNAlater (Invitrogen) and stored at -80°C. Tissues were homogenized using stainless steel beads and a TissueLyser (Qiagen) prior to RNA extraction using the RNeasy Mini Kit. RNA quality was assessed using a Bioanalyzer 2100 (Agilent).

*Sequencing*

RNA samples were prepared for sequencing using poly-dT enrichment (Illumina) according to the manufacturer's protocol. Input RNA quantities were standardized at 500 ng for bulk tumor tissues and cancer cells, and 50 ng for macrophages due to limited cell yields. Libraries were constructed using the NEBNext RNA Library Prep Kit (Illumina). Library quality control was performed using a Fragment Analyzer (Agilent), followed by quantification with the Library Quantification Kit (Illumina). Pooled libraries were sequenced on a NovaSeq 6000 System (Illumina), achieving a minimum depth of 30 million reads per sample. Raw sequencing reads were aligned to the mm10 mouse reference genome using STAR. High-quality alignments were filtered by MAPQ scores >30 using SAMtools. Gene-level counts were generated with featureCounts against GENCODE annotations using the parameters -p -t exon to account for paired-end reads and exon-level quantification.

*Data analysis*

All RNA sequencing analyses were performed in R (v4.3.0). Differential gene expression was assessed using DESeq2 (v1.40.2) with thresholds of $\log_2$ fold change > 0.585 (1.5x linear fold change) and adjusted p-value < 0.05 (Benjamini-Hochberg correction). For functional enrichment analyses, gene ontology (GO) term analyses were conducted using the enrichGO function from clusterProfiler (v4.8.3) with Benjamini-Hochberg adjusted p-value < 0.05. Results were visualized as dot plots (split by ontology for cancer cells) and bar plots (for macrophages) using ggplot2 (v3.5.1). The mouse MSigDB Hallmark gene sets were downloaded, and gene set enrichment analysis (GSEA) was performed using the GSEA function from clusterProfiler (v4.8.3) with default parameters. Statistical significance was determined by false discovery rate (FDR) with a q-value < 0.05, adjusted by the Benjamini-Hochberg method. GSEA results were visualized using the gseaplot2 function from enrichplot (v1.20.3), displaying the normalized enrichment score (NES), enrichment profile, and ranked gene list. Hypoxia-induced pro-angiogenic markers and collagens were visualized as Z-score-normalized heatmaps using pheatmap (v1.0.12), with row-wise scaling of normalized read counts.

**Detailed ECM proteomics**

*Sample preparation*

Tumor tissues were snap-frozen in liquid nitrogen and stored at −80°C until processing. ECM enrichment was performed using the Subcellular Protein Fractionation Kit for Tissues (87790, Thermo Scientific), following the manufacturer’s protocol. Tumor tissues were homogenized in CEB buffer supplemented with 1:100 protease inhibitor cocktail using a TissueLyser (Qiagen). Intracellular protein fractions were discarded, and the final insoluble pellet enriched for ECM proteins was washed three times with PBS containing protease inhibitor cocktail (1:100).

*Mass spectrometry & Protein identification*

All proteomic steps - from protein solubilization and digestion to LC-MS/MS analysis - were performed by the Proteomics Research Infrastructure at the University of Copenhagen. Protein precipitated pellet were lysed in Easypep lysis buffer (A45735, Thermo Fisher) and Pierce Universal Nuclease 100kU (88702, Thermo Fisher) in a 50:1 (v:v) ratio by sonication in a Bioruptor pico (15 cycles, 30s on/off, ultra-low frequency) for 3 times. After, samples were also disrupted using BeatBox (Preomics) with two cycles of 10 min BeatBox Std setting. Samples were incubated for 10 min at 95°C and then, protein concentration was calculated by BCA assay (Pierce). Samples were reduced with 5 mM TCEP for 15 min at 55°C, alkylated with 20 mM CAA for 30 minutes at RT, and digested adding Trypsin/LysC at 1:50 enzyme/protein ratio at 37°C for 16h. Peptides were desalted using EasyPep peptide clean-up plates (A44522, Thermo Fisher) following the manufacturer's instructions. Eluates were dried by vacuum centrifugation, resuspended in buffer A* (2% ACN, 0.1 % TFA) and quantified with Tecan Lunatic. After the quantification, samples were diluted to 0.1 µg/µL in a TwinTec plate for MS analysis. Peptides were separated on an Aurora TS (Gen3) 25 cm, 75 µM ID column packed with C18 beads (1.6 µm) (IonOpticks) using a Vanquish Neo (Thermo Fisher Scientific) UHPLC. Peptide separation was performed using a 50 min stepped gradient of 2-17% solvent B (0.1% formic acid in acetonitrile) for 33 min, 17-25% solvent B for 11 min, 25-35% solvent B for 6 min, using a constant flow rate of 400 nL/min. Column temperature was controlled at 50 °C. Upon elution, peptides were injected via an EASY-Spray source into a Tribrid Ascend mass spectrometer (Thermo Scientific). Spray voltage was set to 1800 . Data was acquired in data independent mode with the Orbitrap MS resolution 60.000 for full scan range 400-900m/z, AGC target was 250% and maximum injection time set to auto. 42 DIA scans with 12 Th width and 1 Th overlap spanning a mass range of 400-900 m/z were acquired at 15,000 MS resolution, AGC target 1000% and maximum injection time of 27 ms. HCD fragmentation normalised collision energy (NCE) was set to 30%. MS files were processed using DIA-NN (v.1.8.2) in directDIA mode with a library predicted from Mus musculus Uniprot fasta file (UP000000589) covering 54,527 protein isoforms. Highly heuristic protein grouping and Match between runs (MBR) were turned on. Carbamidomethylation of cysteine was specified as fixed modification, oxidation of methionine, acetylation at the protein N-terminus, and N-terminal methionine excision were set as variable modifications. Maximum missed cleavage was set to 1 and a maximum of 2 variable modifications were allowed. The minimum peptide amino acid length was 7. The protein groups and precursors were filtered at 1% FDR. All other settings were set as default.

*Data analysis*

Statistical analysis was performed using a developed python code Proteomelit (v1.0.0), based on the automated analysis pipeline of the Clinical Knowledge Graph (1), or using R (v4.3.0). Intensity values were log2-transformed. Uncharacterized proteins and duplicate isoforms were removed. Proteins were retained if they were detected in at least five out of six replicates. Proteins present in four or more replicates in one group and one or fewer replicates in the other were classified as proteins only present in one group. Remaining missing values were imputed using a mixed imputation strategy, where kNN and MinProb methods are used for values missing at random (MAR) and values missing not at random (MNAR), respectively (2). Differentially expressed proteins were identified using unpaired t-tests. *P* values were corrected for multiple hypothesis with permutation-based FDR correction with following parameters: FDR < 0.05, log2 fold-change threshold < 1 and 250 permutations (3). ECM and ECM-associated proteins were annotated using the MatrisomeAnalyzeR tool (v1.0.1) to distinguish them from non-ECM proteins based on annotations from MatrisomeDB, a public protein database for ECM and ECM-associated proteins (4).

**References**


1. Santos A, Colaço AR, Nielsen AB, Niu L, Strauss M, Geyer PE, et al. A knowledge graph to interpret clinical proteomics data. Nat Biotechnol. 2022 Jan 31;40(5):692–702. doi:10.1038/s41587-021-01145-6
2. Lazar C, Gatto L, Ferro M, Bruley C, Burger T. Accounting for the multiple natures of missing values in label-free quantitative proteomics data sets to compare imputation strategies. J Proteome Res. 2016 Apr 1;15(4):1116–25. doi:10.1021/ACS.JPROTEOME.5B00981,

3. Tyanova S, Cox J. Perseus: A bioinformatics platform for integrative analysis of proteomics data in cancer research. Methods Mol Biol. 2018;1711:133–48. doi:10.1007/978-1-4939-7493-1_7,
4. Petrov PB, Considine JM, Izzi V, Naba A. Matrisome AnalyzeR - a suite of tools to annotate and quantify ECM molecules in big datasets across organisms. J Cell Sci. 2023 Sep 1;136(17):261255. doi:10.1242/JCS.261255
